# 3D-MINSTED nanoscopy and protein tracking in densely labelled living cells

**DOI:** 10.64898/2026.03.09.710616

**Authors:** Julius Bernhard Peters, Christopher Heidebrecht, Michael Weber, Marcel Leutenegger, Stefan W. Hell

## Abstract

Investigating the movements and conformational changes of proteins in living cells is essential for understanding their function. The recently introduced fluorescence nanoscopy method called MINSTED has successfully tracked single fluorophore-labelled proteins, albeit in fixed cells and two dimensions only. Here we introduce a MINSTED setup for live-cell tracking of individual proteins in three dimensions (3D) with a localization precision σ down to < 1 nm. Applied to the motor protein kinesin-1, our MINSTED nanoscope follows single proteins in 3D as they process along microtubules in 16 nm steps. Individual steps are resolved amid substantial intracellular background. The unique capability of the 3D stimulated emission depletion (STED) beam to carve out the signal of the protein label of interest enables efficient single-molecule investigations at hitherto unpractically high concentrations of labelled proteins and thus in crowded live-cell environments.

## Introduction

Fluorescence super-resolution microscopy or nanoscopy has evolved into a powerful method to study the distribution and movements of fluorescently labelled biomolecules in cells. The recently developed nanoscopy method called MINFLUX^1,2^ provides single-fluorophore localization with a precision ranging from single-digit nanometers down to Angströms^3^, as well as protein tracking with unprecedented nanometer/millisecond spatiotemporal resolution. Consequently, in a number of pioneering studies, MINFLUX provided new insights in several fields, including the conformational change of the touch-related protein PIEZO-1^4^, the molecular architecture of the photoreceptor’s active zone^5^ or the concurrent diffusion of nicotinic acetylcholine receptors and fluorescent cholesterol^6^. Furthermore, the unique tracking capabilities have shed new light on the stepping of motor proteins, specifically kinesin^7,8^ and dynein^9^, in fixed and living cells.

In a nutshell, MINFLUX establishes the position of a fluorophore by probing its emission rate with a focal excitation intensity distribution featuring a central minimum whose position is precisely controlled by an electrooptic beam steering device. The intensity gradient around the minimum and the concomitantly changing emission rate is exploited to narrow down the difference between the unknown position of the fluorophore and the known position of the minimum, ultimately revealing the fluorophore’s position. To attain a certain localization precision, MINFLUX typically requires about 100 times fewer detected photons than the established method of calculating the centroid of the fluorescence diffraction pattern left by the fluorophore on a camera. Two-dimensional (2D) MINFLUX in the focal plane (x,y) of a microscope is typically performed with a donut-shaped beam^1^ featuring a central minimum that is ideally zero. Localization in three dimensions (3D) is accomplished by a so-called 3D donut or bottle beam^2,3,7,10^ featuring a central minimum confined in all directions (x,y,z) (Supplementary Fig. S1).

Seeking to minimize the fluorescence from the localized fluorophore, MINFLUX requires a background that is persistently even smaller. Unfortunately, due to the laws of diffraction, the 2D and the 3D donut-shaped excitation beams used in MINFLUX cover a much larger sample volume than a regularly focused beam. Besides, donut-shaped beams concentrate most of their power in their periphery, i.e., their crest, that is not used for localization. Thus, donut-shaped excitation beams inherently generate more unspecific background than regularly focused beams. In addition, they produce unwanted fluorescence from neighbouring fluorophores, unless the fluorophores are at least 3-5 times further apart than in standard localization. Concretely, a typical 2D focal donut of 580 nm wavelength features a peak-to-peak distance of about 400 nm in the focal plane and a full width at half maximum (FWHM) of about 500 nm along the optical axis (Supplementary Fig. S1). In a relatively crowded region with many excitable fluorophores, the fluorescence from neighbouring fluorophores easily surpasses that of the fluorophore to be tracked. MINFLUX implementations typically counteract such adverse signals by applying confocal detection with a pinhole that is optically conjugate to the donut minimum. This measure reduces the volume of detection in the sample, creating an effective point-spread-function (E-PSF) that is spatially more confined than the actual volume of excitation. Still, the unfavourable background situation is exacerbated by the fact that during the MINFLUX process of moving the minimum closer to the fluorophore, the signal of the fluorophore decreases with respect to that of other fluorophores and other background. At the end of the day, despite the advancements brought about by MINFLUX localization, tracking with nanometer/millisecond spatiotemporal resolution in fluorophore-rich and crowded cells has remained challenging, if not impossible.

The more recent method called MINSTED^11–13^ features a conceptual advantage over MINFLUX in this regard. Instead of using an excitation minimum, MINSTED uses a regularly focused but pulsed excitation beam that is overlapped and temporally interlaced with a donut-shaped pulsed beam for stimulated emission depletion (STED)^11–13^. The STED beam pulses suppress fluorescence emission throughout the excitation focal volume, except near the intensity minimum of the STED donut^14^. Theory predicts that, at typical settings, the application of a 2D STED-donut yields an E-PSF with a ~45 times more confined sampling volume than confocal 2D-MINFLUX, whereas a 3D STED donut yields a sampling volume that is about ~30 times more confined in MINSTED than the 3D donut sampling volume used in confocal 3D-MINFLUX (Supplementary Fig. S2a-d). Motivated by these insights, we now introduce a setup and method for 3D localizing and tracking individual fluorophores by MINSTED. The background suppression provided by this setup tracks single fluorophore-labelled proteins in cells at ~20 times higher densities of labelled proteins than commercial MINFLUX tracking systems, yet at a spatiotemporal resolution that is comparable to 3D-MINFLUX (Supplementary Fig. 2e,f). This unique feature is demonstrated by recording the stepping of single kinesin-1 motor proteins along and around the microtubules in fixed and living cells. A major benefit of being able to track amid many labelled proteins is that the number of succesful measurements per unit time increases accordingly, which greatly improves the applicability of single-molecule 3D tracking in living cells.

## Results

### 3D-MINSTED principle

To enable precise 3D localization, we constructed a beam-scanning confocal fluorescence microscope that, besides providing lateral (x,y) MINSTED localization^11^, features an additional STED beam path for localization in the axial direction (Fig. 1a and Supplementary Fig. S3). All the MINSTED laser beams used in this setup are pulsed, with each 200 ps pulse of excitation light instantly followed by a 1.5 ns pulse for STED. The excitation and STED wavelengths are 532 nm and 636 nm, respectively. The excitation laser is operated at a repetition rate of 30 MHz, while three distinct STED lasers (STED XY, STED Z1 and STED Z2), operating at a lower repetition rate of 10 MHz, probe the lateral and axial position in an interleaved fashion.

**Fig. 1.**
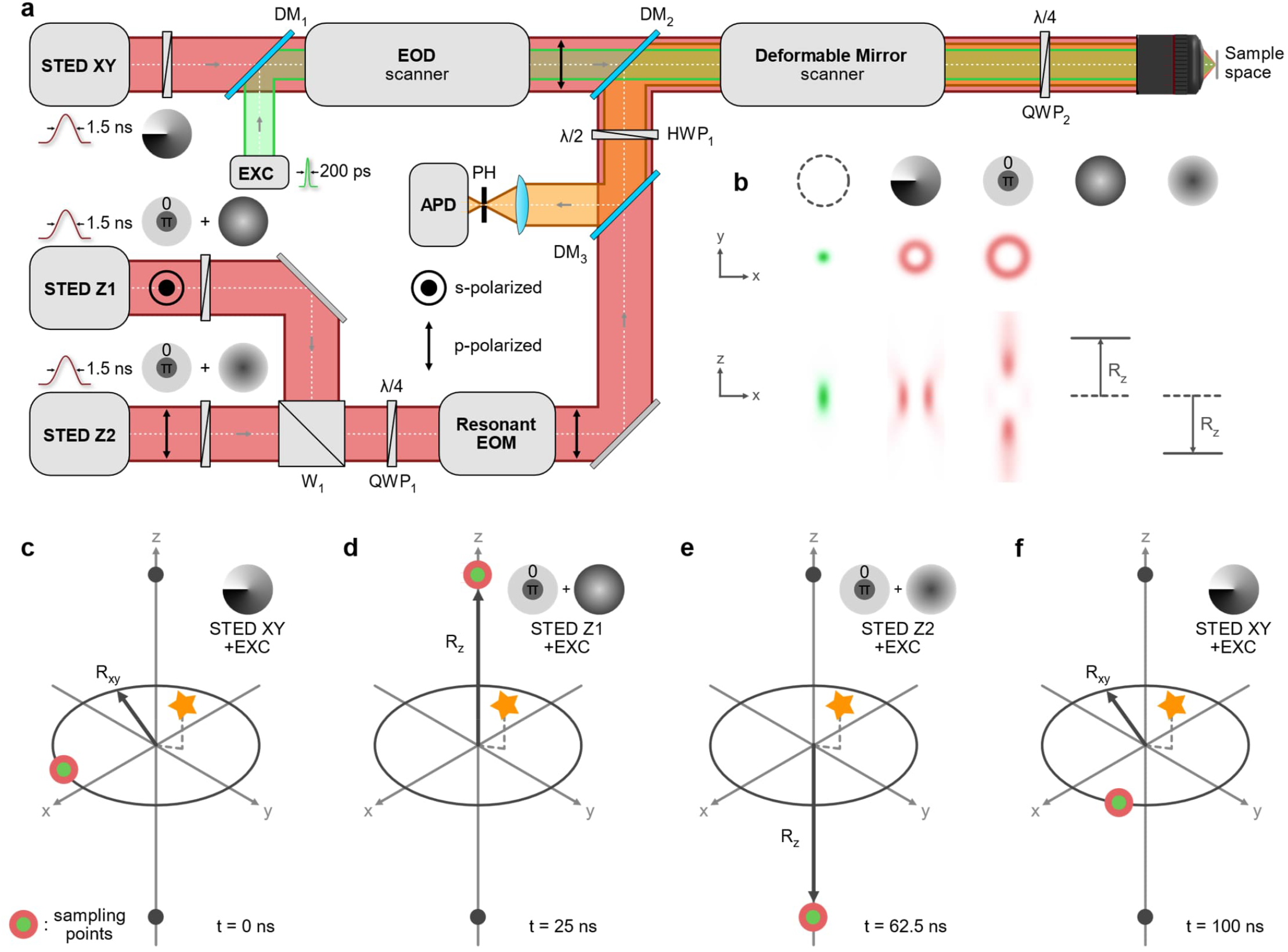
MINSTED localization in three dimensions. **a** Schematic of the 3D-MINSTED setup: the lateral localization relies on 1.5 ns long STED pulses (STED XY) that are converted into a donut shape by a vortex phase plate and are aligned with a 532 nm excitation laser emitting 200 ps short pulses. These co-aligned beams are steered in the lateral focal plane of the objective by an EOD scanner, circling the beams in the lateral plane. For axial localization, two additional STED beams (STED Z1/Z2) are converted into 3D donuts using top-hat phase masks and joined by polarization using a Wollaston prism (W_1_). Applying a negative defocus on one of the beams and a positive defocus on its counterpart shifts the two beams to positions ±*R*_z_ above and below the focal plane. The polarization is shifted to p-polarization for both beams with the quarter-wave plate (QWP_1_) and resonant EOM. All beams are combined by a polarization sensitive dichroic mirror (DM_2_). The deformable mirror controls the focus position of those beams by modulating the wavefront, whereas the quarter-wave plate (QWP_2_) ensures circular polarization as required for the XY donut. The fluorescence is extracted by the second polarization sensitive dichroic mirror (DM_3_), spatially filtered by the pinhole (PH) and detected by the APD. **b** A Gaussian excitation spot and the STED foci created by the different phase masks are shown at the focal plane of the objective. **c-f** Repetitive pulse sequence used to sample a fluorophore (represented as a star). The current positions of the STED pulses with the applied wavefront shift and the corresponding excitation pulse at a given time are outlined, where 2*R*_*i*_ corresponds to the FWHM of the E-PSF at the maximum energy of the STED pulses.

For lateral localization, the nanosecond STED pulses are provided by the STED-XY laser (Fig. 1a). Initially Gaussian, the STED-XY beam is converted into a donut beam using a vortex phase plate (Fig. 1b). During the MINSTED localization process, the co-aligned beams are steered along a circular path around the assumed fluorophore position in the focal plane (x,y) of the objective lens by electro-optic deflectors (EODs); see Fig. 1c for a single excitation pulse and a STED pulse in the lateral plane at *t* = 0 ns. STED confines the spatial region from which a signal photon can originate to sub-diffraction volumes^14^. Previous works showed that photons detected from the tail of the E-PSF carry more information about the fluorophore position than photons detected near the maximum of the E-PSF^11^. On the other hand, the emission rate is lower and the bleaching rate higher when collecting photons at the tail^1,11,15^. For the right balance between spatial information gain per single detected photon and the rate of emitted photons, the circle radius *R*_*xy*_ was chosen to be half of the full width at half maximum (FWHM) of the E-PSF in the lateral plane^11^.

For axial MINSTED localization, the same principle of fluorescence confinement is implemented accordingly. Instead of sampling along a continuous circular path, two additional STED laser beams are utilised to sample above (STED Z1) and below (STED Z2) the estimated fluorophore position at a distance ±*R*_z_ from the focal plane (Fig. 1d and 1e at different times *t*). The wavefronts of the STED Z beams are imprinted with a tophat phase profile including the defocus, using a spatial light modulator (SLM), such that appropriately shifted 3D donuts are created (Fig. 1b). The distances ±*R*_z_ were chosen to be about half the FWHM of the E-PSF at maximum STED pulse energy in the axial direction (Methods). Due to the comparatively slow update rate of the SLM of typically 60 Hz, *R*_z_ was kept constant during a localization. The centre position along the axial direction lies in the plane for the lateral MINSTED localization (Fig. 1d,e). The STED Z beams, which are mutually orthogonally polarized, are joined by a Wollaston prism (W_1_), and finally transmitted through an achromatic quarter-wave plate (QWP_1_) and a resonant electro-optical modulator (Res-EOM). The combination of quarter-wave plate and Res-EOM, driven at a frequency of 20 MHz, periodically converts s- to p-polarization, shifting the orthogonally polarized sequential pulses into a single p-polarized pulse stream. Since the p-polarized light is always transmitted at the polarization sensitive dichroic mirrors (DM2, DM3) the single pulse stream is converted to s-polarization by the achromatic half-wave plate (HWP_1_). Thus, all beams are combined at DM_2_, which reflects and transmits the s-polarized STED Z light and the p-polarized STED XY light, respectively. This arrangement enables the three STED laser beams of the same wavelength to be combined with minimal losses.

The position of the specific donut with their corresponding excitation pulse defines the confinement of the E-PSF on the circular path for STED XY, at the upper position for STED Z1 and at the lower position for STED Z2, each one independently. At the same time, the position of the donut minimum provides a perfectly defined reference coordinate in the sample. By modulating the wavefront, the deformable mirror controls the focal point of all beams and sets field of view (FOV); it also descans the fluorescence signal. The second quarter-wave plate (QWP_2_) ensures circular polarization. Upon detection of a photon by the avalanche photo diode (APD), the common focal point of all beams is updated towards the most recent position of the STED donut that defined the E-PSF causing the photon detection. Thereby, the common focal point ultimately circulates or wiggles around the fluorophore position until the fluorophore is bleached or lost^11^.

### Imaging of 2D structures in 3D space

To demonstrate 3D-MINSTED nanoscopy and its localization performance, planar 3×3 DNA origami grids were bound to the coverslip by a biotinylated bovine serum albumin streptavidin complex and measured in solution with DNA-PAINT^16^. The DNA origami grid constants were specified by the manufacturer GATTAQuant to be 10.9 nm parallel to the DNA helices and 12 nm orthogonal to them. So-called imager strands, i.e. single-stranded DNA labelled with one Cy3B fluorophore, docked stochastically to any of the nine potential binding sites of the grid, while other fluorescent imager strands diffused freely in the solution (Fig. 2a). By applying the described 3D-MINSTED procedure, the docked fluorophores were localized, resulting in about 600 localizations of the binding sites on 87 automatically detected grids in 7 different FOV (Methods). For each detected grid, a 3×3 grid, representing the expected binding sites was matched to the localizations. Thus, we obtained the grid constants *d*_x_ and *d*_y_. An example of a flat lying DNA-origami with eight measured binding sites and the corresponding grid constants are displayed in the lateral projection (Fig. 2b). The coloured circles show the average measured positions attributed to the nearest binding site of the matched grid (grey dots). In the same field of view, line-shaped objects in the lateral projection can be observed (Fig. 2c). After 3D rotation into a lateral plane, the grid structure clearly reveals itself (Fig. 2d). The arbitrary rotation of grids may be caused by unspecific binding of biotin to one of the four possible binding sites of streptavidin^17^ and by the 3D topography of the fixating layer.

**Fig. 2.**
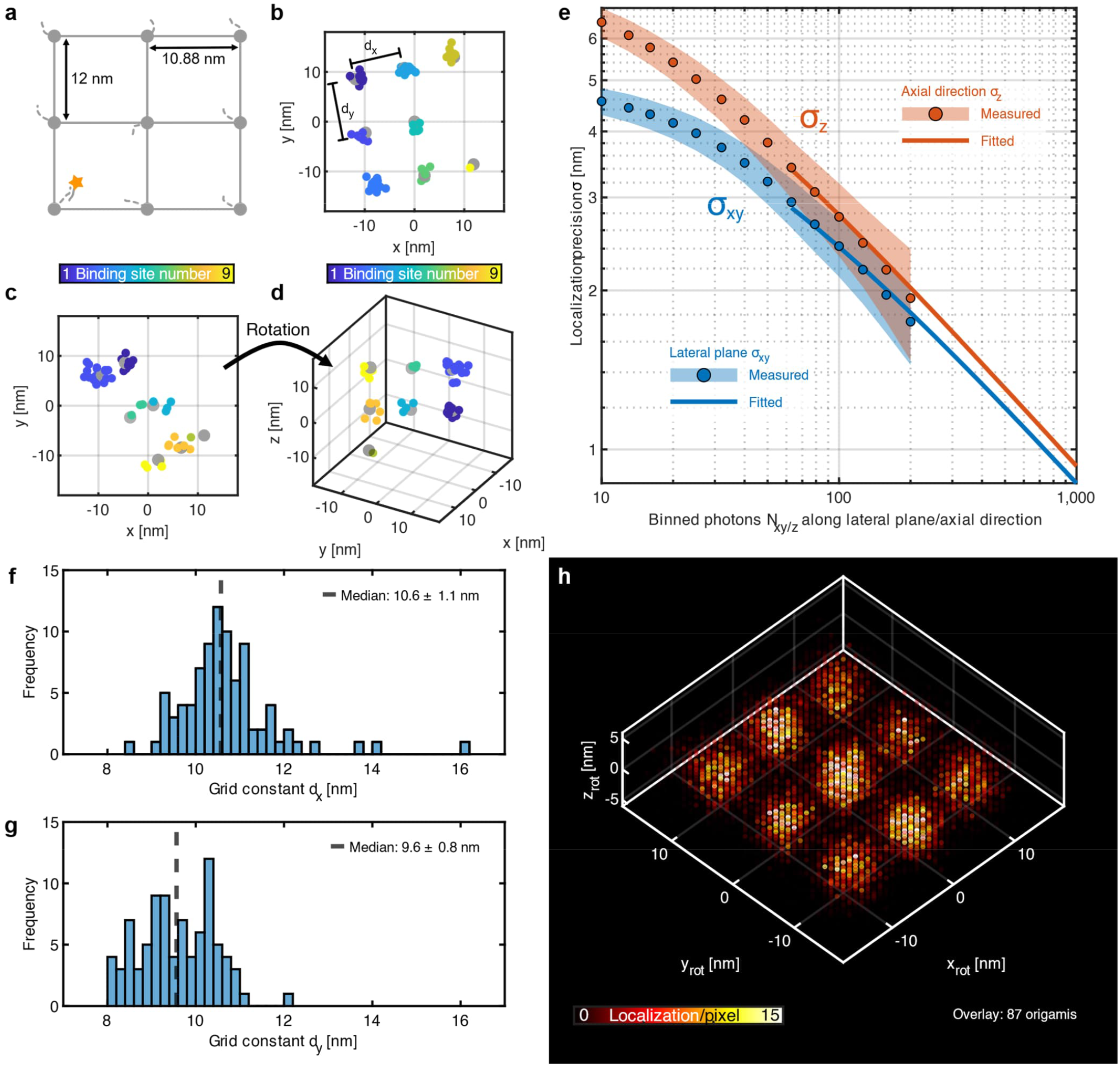
3D-MINSTED nanoscopy with 2D samples tilted in 3D. **a** Schematic of a 3×3 DNA origami grid with nominal grid constants of 10.9 nm and 12 nm, respectively, measured with DNA-PAINT. The Cy3B fluorophore is represented with the star attached to an imager strand. **b** A flat-lying DNA origami grid in the lateral plane with eight measured binding sites and the extracted grid constants *d*_x_ and *d*_y_ is shown. **c**,**d** A rotated DNA origami grid in the lateral plane (**c**) and rotated in 3D (**d**) with seven binding sites is depicted. The coloured circles are the average localization positions attributed to the nearest expected binding site. Grey dots mark the expected binding sites of the matched regular 3×3 grid. **e** Localization precision of all evaluated localizations (points: median; shaded areas: ±1 standard deviations) along the lateral plane σ_xy_ (blue) and the axial direction σ_z_ (red) depending of the binned number of photons *N*_*xy*/z_ in the corresponding direction. The localization precision of an individual localization is estimated by a fit (solid lines). **f**,**g** Measured grid constants along the rotated x-axis x_rot_ (**f**) and y-axis y_rot_ (**g**) with the corresponding mean values (dashed line). The grids were rotated such that *d*_x_ > *d*_y_. **h** Overlay of all evaluated estimated localization positions from 87 identified origamis (Supplementary Fig. S8) with 200 photons binned along the corresponding axis in a 4D histogram, resolving the 3×3 DNA origami. Each circle represents the number of localizations in a 0.5 nm × 0.5 nm × 0.5 nm volume.

Due to the separate lateral and axial position updates of the 3D-MINSTED localization, the pertinent localization precisions can also be treated separately. First, the centre positions of a localization track were split into two sequences of lateral and axial position updates, respectively. The standard deviations 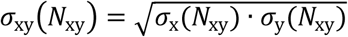 and σ_z_(*N*_z_) for bins of *N*_*xy*_ and *N*_z_ photons, were used to estimate the localization precisions. For this purpose, the centre positions were averaged for each bin and the standard deviations for each bin length were calculated as long as at least 5 non-overlapping bins were available. To estimate the localization precisions of a complete track, these *N*-photon localization precisions were fitted to a polynomial model σ_fit_(*N*) = *a*/(*b* + *N*)^*c*^ with 0 < *a, b* and 0 < *c* < 0.5 as presented in previous works^12,13^. Thus, we achieved localization precisions down to < 1 nm in all directions (Fig. 2e), which in turn enabled us to localize and attribute individual binding sites of the DNA origami grid with high fidelity.

For further analysis, the identified origami grids were rotated into a common plane, such that all grids were mutually aligned by their expected binding sites. We oriented the larger grid spacing along the rotated x-axis x_rot_ and the shorter grid spacing along the rotated y-axis y_rot_ (Supplementary Fig. S8). Thereby, we measured median grid constants of *d*_x_ = (10.6 ± 1.1) nm and *d*_y_ = (9.6 ± 0.8) nm, respectively, confirming that 3D-MINSTED can easily resolve these structures.

The manufacturer GATTAQuant specified the DNA origami grid constants to be 10.9 nm and 12 nm. The first grid spacing extends along the DNA helices and is expected to be stiff. It corresponds closely to our measured *d*_x_^18^. The linker of six carbon atoms from the strand to the fluorophore with a length of about 1 nm introduces some uncertainty by allowing the fluorophore to move around the binding site^19^. The origami shape varies with salt-concentration by up to 20% due to a rolling-up of the rectangular DNA stacks^20^. Our measured *d*_y_ indicates moderate shrinkage as expected at the 75 mM magnesium chloride salt-concentration of the imaging buffer (Methods). A visual confirmation of resolving the structure is depicted in Fig. 2h, where all localizations attributed to the 87 aligned origamis are shown in a 4D histogram for 200 binned photons each in the lateral and axial directions. The cluster precision as a function of binned photon number is also characterized (Supplementary Fig. S6).

### Tracking in fixed cells

The cytoskeletal motor protein kinesin-1, hereafter referred to as kinesin, plays a key role in intracellular transport along microtubules. Kinesin transports cargos towards the plus end of microtubules under ATP consumption^21,22^. Its processive movement follows a hand-over-hand walking mechanism, in which the trailing head overtakes the microtubule-bound leading head, thereby becoming the new leading head. This continuous sequential binding and unbinding of the microtubule leads to observable positions where kinesin binds the microtubule. For a kinesin labelled at the motor domain (head) these centre positions are expected to be 16 nm apart (Fig. 3b). In fact, MINSTED was already shown to track kinesin molecules^13^, albeit only on artificial microtubules synthesized from purified tubulin monomers. Also, MINSTED tracking was not accomplished in cells, neither fixed nor living, and just in 2D. Hence, a major motivation for this study was to find out if and to which extent the unique fluorescence suppression brought about by STED bestows 3D-MINSTED with the ability to track single proteins in fixed and living cells.

**Fig. 3.**
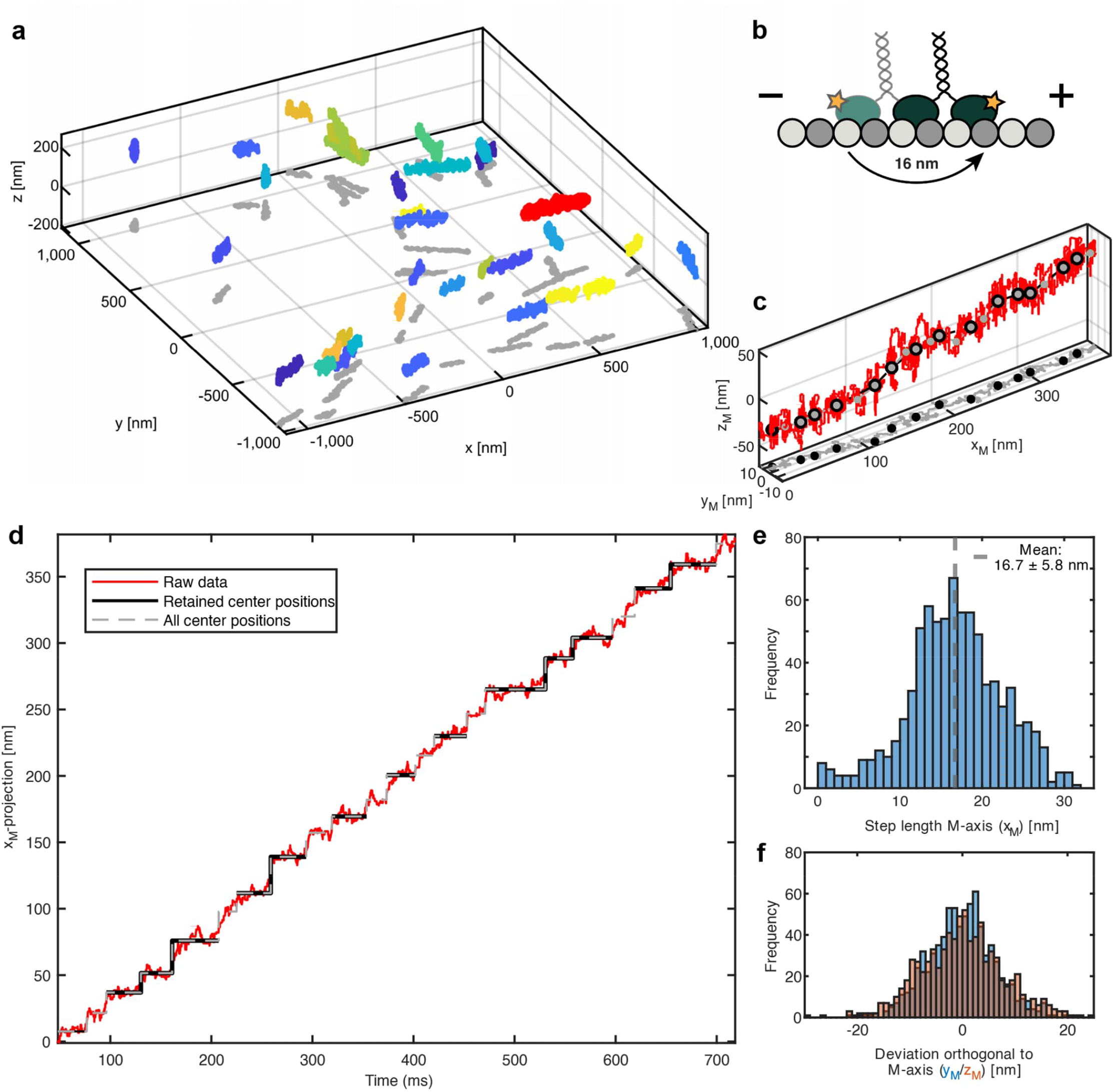
Kinesin tracking in fixed cells by 3D-MINSTED. **a** Overview of 3D tracks of kinesin labelled at the motor domain observed in fixed U-2 OS cells. Separate tracks are displayed in separate colours with their projection onto the xy plane (grey). **b** Schematic representation of kinesin’s hand-over-hand walking mechanism on microtubules. The expected step size with a fluorophore attached to the motor domain is approximately 16 nm. **c** Projection of an exemplary track (highlighted in **a** in red). Raw localizations (red line), with fitted centre positions (grey filled dots), retained steps (black circles), along with their projection onto the xy plane. **d** Position versus time plot of projection of the highlighted track. Fits of the steps (gray dashed line) retained for further analysis (black line). **e** Step length histogram of 788 retained steps from all 188 retained tracks with mean step length of (16.7 ± 5.8) nm along the microtubules. **f** Histogram of distances from the microtubules (orthogonal deviation from the x_M_-axis) in y_M_ (blue) and z_M_ (orange).

To this end, we first labelled individual kinesin molecules at the motor domain in fixed U-2 OS cells at 500 µM ATP concentration (Fig. 3a) with the fluorophore Cy3B and recorded its translation in space. To maintain the 3D architecture of microtubules, we used a gentle fixation protocol that we adapted from the MINFLUX kinesin tracking study conducted by the Ries group^7^. We then added an imaging buffer containing motor-domain labelled kinesin and ATP. Direct labelling of microtubules was not required to identify cells, because we were able to densely decorate microtubules with the labelled kinesin. The kinesin tracks were selected automatically. From a total of 1440 measured tracks, the selection algorithm provided us with 188 tracks for further analysis (Methods). By estimating the microtubule shape with a third order polynomial, these tracks were analysed along a projected microtubule axis (x_M_-axis) (Methods). We observed on average 16 nm steps both from 3D renderings of individual tracks and from projected x_M_(t) plots (Fig. 3c,d). The ATP concentration of 500 µM was comparable to physiological ATP concentrations. The high speed of kinesin at this ATP concentration often resulted in a step of the kinesin before the MINSTED algorithm converged to the previous kinesin position. Therefore, we excluded steps where we could not confidently estimate the centre positions before and after a step. Furthermore, we performed the step length analysis only on tracks containing at least two confident steps.

Altogether, we retained 188 kinesin tracks with a total of 788 kinesin steps. Fig. 3e shows a histogram of the extracted step lengths. From all extracted steps, we obtained a mean step length of (16.7 ± 5.8) nm along the microtubule (Fig. 3e), confirming previous 2D tracking measurements of kinesin by MINSTED^13^ and MINFLUX^7,8^. We further analysed the tracks perpendicular to the x_M_-axis in the lateral plane (y_M_-axis) and along the optical axis (z_M_-axis) (Fig. 3f). In both y_M_- and z_M_-directions, we observed excursions of up to about 20 nm. Whereas systematic deviations could arise from kinesin sidestepping to different protofilaments or even microtubules^7,13^, random excursions are most likely caused by the single-photon updates of the estimated fluorophore position during the MINSTED localization. As both histograms follow zero-mean normal distributions, we attribute the excursions to the localization process.

### Tracking in living cells

Finally, we tracked kinesin in living cells and hence in a high-background environment that poses major challenges for 3D tracking of single proteins otherwise. For this purpose, we adapted the protocol for live-cell kinesin tracking^7^. We applied an overexpressing kinesin plasmid with a HaloTag at its motor domain to transfect U-2 OS cells. Afterwards, we labelled with a concentration of 100 nM JF549 HaloTag and after washing obtained cells with a labelling density entailing more than a single fluorophore within the diffraction-limited sampling volume (Fig 4a). For all commercially available super-resolution localization techniques and especially MINFLUX tracking, it is crucial that within the sampling volume, only a single fluorophore is active at a time, because photons arising from two or more emitters clearly invalidate the localization. As a matter of fact, the need for sparse labelling presents a major challenge in living cells where labelling densities are inherently difficult to control.

**Fig. 4.**
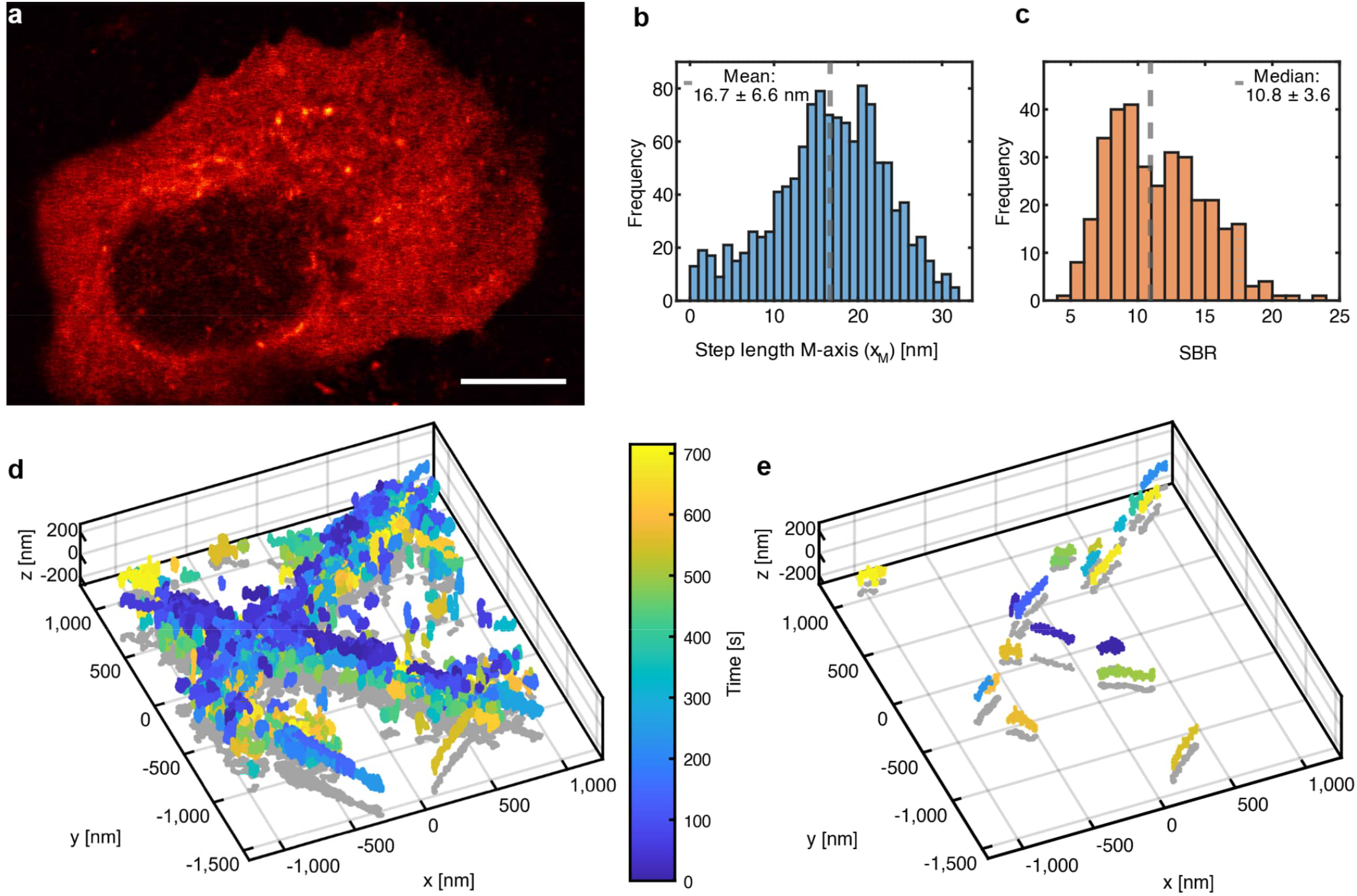
Kinesin tracking in live cells by 3D-MINSTED. **a** Confocal scan of an exemplary U-2 OS cell overexpressing kinesin with a HaloTag on its motor domain. Scalebar: 10 µm. **b** Step length histogram of retained live-cell steps along the projected x_M_-axis. A mean step length of (16.7 ± 6.6) nm (dashed grey line) was measured from 1207 retained steps in 337 tracks. **c** Histogram of SBRs of retained tracks. The median SBR was 10.8 ± 3.6 (dashed grey line). Overview of 3D kinesin tracks, all recorded tracks (**d**) and the filtered single fluorophore track (**e**) in an example FOV are shown.

Although the fluorophore density in the cells exceeded the typical single-fluorophore regime, we efficiently acquired kinesin tracks (> 10000) in 3D on four different cells, each on a different coverslip, with 25 FOVs in total. As in the previous section, the single fluorophore kinesin tracks were selected automatically (Methods). From 337 valid tracks, we blindly extracted a total of 1207 steps for further analysis (Fig. 4b) using the previously described algorithm (Methods). We measured an average step length of (16.7 ± 6.6) nm along the x_M_-axis of the kinesin motor domain, which is in good agreement with the step length obtained by 3D MINFLUX^7^. The potential of MINSTED to supress the background becomes evident from the signal-to-background ratio (SBR) of all retained tracks (Fig. 4c). Even in this densely labeled region we estimated a median SBR of 10.8 ± 3.6 for the live-cell kinesin tracks (Supplementary Fig. S15).

A typical example FOV with all measured tracks (Fig. 4d) reflects the high success rate of single kinesin tracking. The lateral focus-lock of the setup was not activated for these experiments, slightly broadening the width of the observed features due to long-term drift over the measurement time of > 10 min per FOV (Methods). The width of a single ‘branch’, ~200 nm in the overlayed tracks, does not necessarily reflect the shape of a single microtubule. However, because the longest measured time for a track is only 2.4 s, on a single track this long-term drift can be neglected (Supplementary Video 1). This particular FOV yielded 23 automatically selected and valid single fluorophore tracks (Fig. 4e). In total, several hundreds of valid tracks over a single measurement day were recorded, showing the potential of 3D-MINSTED to measure protein dynamics routinely and with high efficiency.

## Discussion

We demonstrated that the super-resolution concept of MINSTED imaging and localization can be effectively realized in 3D, opening up single fluorophore 3D tracking with a precision down to < 1 nm. Compared to 3D-MINFLUX, 3D-MINSTED owes its success primarily to the fact that the excitation beam is overlapped with a 3D-STED beam that provides a 3D reference coordinate in the sample through its nearly zero-intensity point, while also confining fluorescence emission to volumes far below the diffraction limit. Thus, the 3D-STED beam substantially reduces the volume around the tracked fluorophore that must remain devoid of other active fluorophores.

2D fluorophore arrangements are resolved in 3D with an inter-fluorophore distance down to < 10 nm, irrespective of their orientation in space. The deviations from the specified value of 12 nm along the rotated y-axis y_rot_ are probably caused by rolling-up of DNA helices due to elevated magnesium chloride salt-concentrations^20^. Our analysis predicts a localization precision < 1 nm in a given direction provided that 1000 photons are detected to that end. Thus, MINSTED reaches virtually the same localization precision as MINFLUX, but with a higher SBR (Supplementary Fig. S14a).

Tracking the motor protein kinesin-1 in 3D in fixed and living cells revealed the 16 nm steps of a single motor domain with a precision of 4.8 nm, 4.4 nm, 11.7 nm in the x, y and z direction, respectively, and a temporal resolution of 2.4 ms. The tracking was accomplished even in high background in cells, yet the tracks could be evaluated automatically. The reliability of the tracking data was mainly limited by the localization precision along the optical axis (Supplementary Fig. S14 a, c), which is about twice as large as its counterpart in the focal plane. This difference stems from the fact that in the lateral direction the E-PSF was sharpened down to a FWHM of ~36 nm, whereas along the optical axis, a FWHM of only ~80 nm was obtained. The lower precision along the optical axis was due to the limited pulse energy of *E*_z_ ≈ 0.4 nJ provided by the two STED lasers confining the E-PSF in the z-direction. In the future, this disparity can be mitigated by using lasers with a higher pulse energy. A FWHM of ~40 nm of the E-PSF should be readily possible in living cells without inducing adverse effects on the cellular process to be studied. We also note that our optical implementation of 3D-MINSTED can be improved further by resorting to a scheme using a single STED laser for all directions. Such optical improvements will be appealing since our study clearly proves the unique capability of 3D-MINSTED to track individual proteins in densely labelled regions, where tracking has so far failed. By inherently reducing the demands on sample preparation and providing hundreds of single-fluorophore tracks per measurement day, 3D-MINSTED is destined for single-protein tracking in applications that were hitherto unviable.

## Acknowledgement

Miroslaw Tarnawski and the Protein Core Facility at the MPI for Medical Research for the purification of the K28C kinesin construct. Ellen Rothermel and Tanja Koenen for their help with cell culture. The electronic and mechanical workshop of the MPI for Multidisciplinary Sciences under the management of Frank Meyer, Mario Lengauer and Christian Klaba for helping us with custom parts in the setup. Steffen J. Sahl and Henrik von der Emde for the fruitful discussions and critical reading.

## Author contributions

J. B. P. built the setup under the supervision and design inputs of M. W.. M. L. wrote the software, including the real-time control of the setup and implemented the deformable mirror calibration. M. W. and J. B. P. built the fibre-amplified STED laser system after planning and construction by M. W.. J. B. P. prepared the 3×3 grid DNA-origami samples and C. H. prepared and labelled the kinesin-1 on microtubules for fixed and living cells. J. B. P. performed the measurements. J. B. P. evaluated and analysed the data with the help of M. L. and feedback from S. W. H. and C. H.. S. W. H. outlined the research project, initiated and supervised its exploration. S. W. H., M. L., C. H. and J. B. P. wrote the manuscript. All authors contributed to the manuscript and the supplementary information either through discussions or directly. This work has been funded by the Bundesministerium für Forschung, Technologie und Raumfahrt (FKZ 13N14122 to S. W. H).

## Material and Methods

### Experimental setup

The experimental setup is sketched in Supplementary Fig. S3 and its components are listed in the Supplementary Information. The axial localization of a fluorophore reported here necessitated a considerable expansion of the MINSTED setup over previous 2D-MINSTED implementations^11^, involving additional STED lasers and positioning systems. In our setup, the lateral MINSTED localization is realized by overlapping the beam of a ps-pulsed excitation laser emitting at a wavelength of 532 nm with the beam of a self-built ns-pulsed fibre amplified STED laser emitting at 636 nm (636_1_); by deflecting those beams in the pupil planes with two electro-optical deflectors (EOD_x,y_); and by circling their foci around the estimated position of the fluorophore at 125 kHz frequency with a maximum radius of 125 nm. The STED beam passes through a vortex-phase mask in the pupil plane to create the STED donut in the focal plane for lateral emission confinement. After passing the EODs, both beams are p-polarized and reflected by a deformable mirror. By modulating the wavefront of the beams^23^, the deformable mirror is used to scan a field of up to 3 µm × 3 µm × 3 µm extent and shift the centre position in 3D during the localization process. An achromatic quarter-wave plate (λ/4) establishes circular polarization for all beams before being focused into the sample by a silicon oil immersion objective lens of 1.3 numerical aperture (NA). The fluorescence signal is descanned by the deformable mirror, reflected at the polarization-sensitive dichroic mirrors (DM_2,3_) and is detected with a single-photon avalanche diode (APD_1_) at 550–610 nm wavelength.

For confinement of fluorescence in the axial direction, two additional ns-pulsed STED beams at 636 nm wavelength with mutually orthogonal polarization (636_2,3_) pass through a segmented achromatic half-wave plate (λ/2_s_) ensuring s-polarization on the spatial light modulator (SLM). The SLM controls the position and shape of the foci of these additional beams. Phase profiles ensuring a top-hat phase modification and an axial defocus create 3D donuts above and below the focal plane. The SLM allows for aberration correction in the sample^24^. The calibration of the defocus phase profile is described in the Supplementary information. The beams for 3D donut formation are joined with a Wollaston prism (W_1_). The common beam is transmitted through an achromatic quarter-wave plate, a resonant electro-optical modulator (Res-EOM), driven at a frequency of 20 MHz, and a polarizing beam splitter (PBS) to clean up the polarization. The Res-EOM periodically rotates s-polarized light to p-polarization, thereby allowing to combine alternating pulses of the orthogonally polarized beams into one p-polarized beam. Afterwards, the beam transmits through DM_3_, is rotated to s-polarization by an achromatic half-wave plate (λ/2) and is reflected by DM_2_ onto the deformable mirror.

The power of each pulsed laser is controlled by acousto-optical modulators (AOM). The timing of the pulses for the lasers and the analogue sinusoidal wave for the Res-EOM are defined by a static pattern, provided by an arbitrary wave generator.

The setup is based on confocal beam scanning, which can be coupled into the system with a flip mirror. The confocal scanner addresses a field of 64 µm × 64 µm in the lateral plane of the sample with two galvo mirrors^12^. Continuous-wave lasers provide excitation light at a wavelength of 488 nm and 560 nm with a fixed power of around 15 µW in the aperture of the objective lens. The detection part of the setup splits the emitted fluorescence into two colour channels of 495–520 nm and 580–610 nm wavelength ranges.

For calibrating the defocus phase profile (Supplementary Fig. S5), a pellicle is manually placed into the beam in order to deflect a part of outcoming light onto a photomultiplier tube (PMT). Due to the different polarization of the STED beams, the PBS can be rotated to select the corresponding polarization.

To select the region of interest, the sample is mounted on a piezo stage for fine movements. The piezo stage is mounted on three linear stages for coarse positioning. The sample can be actively stabilized both laterally and axially. The axial feedback signal is obtained by detecting the reflected spot of a power-stabilized super-luminescent light emitting diode (SLED) at 1000 nm wavelength from the coverslip sample onto the camera CAM_2_. The lateral feedback signal is obtained by tracking fiducial markers in darkfield images on the camera CAM_1_. The fiducial markers are imaged in a field of 70 µm × 70 µm. They are placed off-axis to avoid interactions with the MINSTED illumination of the sample. For imaging the fiducial markers, the reflected SLED illumination beam is blocked by an iris diaphragm. The fundamental optical structure and the software of the focus locks is described in previous work^12,25^.

The microscope is controlled by two field-programmable gate array (FPGA) devices and a custom graphical user interface implemented in LabVIEW 2021 (National Instruments) and MATLAB R2021b (The Mathworks, Natick, MA, USA).

### MINSTED measurements

A fluorescent molecule was searched by applying a confocal scan in a lateral region of interest (ROI) using the deformable mirror. When at least *N*_on_ photons were detected in a 2 by 2 pixels neighbourhood, a 3D-MINSTED localization was initiated^11^.

The lateral localizations were executed with a sampling pattern radius *R*_xy_ equal to half the FWHM of the lateral E-PSF. Upon each lateral photon detection, the lateral centre (*x, y*) of the pattern was moved *α*_*xy*_*R*_xy_ towards the causative sampling position; the STED energy was increased such that the FWHM of the lateral E-PSF was reduced to the radius *γ*_xy_*R*_xy_ down to a preset minimum. We used *α*_xy_ = 15% and *γ*_xy_ = 97% as established previously^11–13,23^. The axial localizations were performed with a fixed axial sampling at ±*R*_z,samp_ equal to 0.75 *R*_z_ (for imaging) and 0.5 *R*_z_ (for tracking), where *R*_*z*_ is half of the minimal FWHM of the axial E-PSF. Upon each axial photon detection, the axial centre *z* of the sampling pattern was moved *α*_z_*R*_z_ towards the causative sampling position and the FWHM of the axial E-PSFs was reduced to *γ*_z_*R*_z_ down to a preset minimum. Supplementary Fig. S4 shows the correlation between *α*_z_ and the simulated localization precision in the axial direction σ_z_. We typically applied *α*_z_ = 10% and a fraction *γ*_z_ = 97%, which proved robust at low SBR. The minimal STED pulse energies were *E*_xy_ = 0 nJ and *E*_z_ = 0.01 nJ and corresponded to minimal radii *R*_xy_ = 118 nm and *R*_z_ = 200 nm. Maximal pulse energies of *E*_*xy*_ = 0.62 nJ and *E*_z_ = 0.4 nJ defined a scan radius of *R*_xy_ ≥ 18 nm in the lateral plane and *R*_z_ ≈ 40 nm along the axial direction. Both lateral and axial localizations were performed simultaneously by cycling through the three STED beams pulse by pulse. Each excitation and STED pulse pair was placed in a separate interval to allow for an unambiguous attribution of the detected photons (Fig. 1c-f). The 3D-MINSTED localization was terminated if less than *N*_off_ = 16 photons were detected during an interval time *t*_off_ = 8 ms.

The DNA origamis samples, ordered from the manufacturer GATTAQuant, were prepared with an origami dilution of 1:9 in 10 mM MgCL_2_ in PBS on coverslips. Gold beads of 200 nm diameter (EM.GC200/7, BBI Solutions) were placed on and served as fiducial markers on the coverslip for the lateral focus lock. The origamis were imaged with DNA-PAINT using a Cy3B-labelled imager strand concentration of 7.5 nM. After five minutes of incubation time, the unbound origamis were washed out four times with 75 mM MgCl_2_ in PBS. Afterwards, the sample was sealed in a solution of 200 μL of oxygen-deprived reducing-oxidizing buffer with a MgCl_2_ concentration of 75 mM^26^. A more detailed description of the sample preparation is described in the literature^12^. To ensure that the DNA-PAINT strands were bound to the DNA origami grid, a search dwell time of 1.6 ms per 80 nm pixel and a threshold of *N*_ON_ = 40 photons were chosen. The sample was actively stabilized in all directions, because the measurement time for a single FOV was around 40 minutes.

For the tracking experiments the search was performed with a dwell time of 60 µs per 75 nm pixel. The threshold was set to *N*_ON_ = 2 photons to measure as many potential tracks as possible for a-posteriori filtering during the data analysis. Measurements were performed on four different cells (Supplementary Fig. S11) with different FOVs per cell. Here, gold beads could not be used as fiducial markers due to their incompatibility with cells. Hence, only the axial direction was actively stabilized. For a single track no significant drift was expected since the maximum measured time for a single track was 2.4 s.

### Kinesin labelling

Cysteine-light truncated (at amino acid position 560) human kinesin-1 with a single solvent-exposed cysteine at amino acid position 28 were expressed in E. coli using the plasmids K560CLM K28C that were purified as described in a previous study^8^. Kinesin-1 was labelled with Cy3B maleimide (19380, Lumiprobe) overnight at 4° C. Excess dye was removed using ZEBA spin desalting columns 7K MWCO (89877, Thermo Fisher Scientific). The concentration and the degree of labelling efficiency was measured using UV-vis spectroscopy and mass spectrometry. Labelled kinesin was supplement with 10 % (w/v) sucrose and stored at −80° C until further use.

### Cell culture

U-2 OS cells were cultured in McCoys media (16600082, ThermoFisher Scientific) supplemented with FBS Superior (S 0615, Bio&SELL), Pen/ Strep, (P0781, Sigma Aldrich), Natirum-Pyruvat (S8636, Sigma Aldrich) and MEM NEAA (M7145-100ML, Sigma Aldrich). The cells were kept at 37° C, 5 % CO_2_ and 100 % humidity.

### Fixed-cell kinesin tracking

U-2 OS cells were seeded at least one day prior to experiments on 18 mm round microscopy coverslips. For fixation, the cells were incubated in extraction buffer (PEM80: 80 mM PIPES, 1 mM EGTA, 1 mM MgCl_2_ set to pH 7.4 supplemented with 1 mM sucrose and 0.15 % Triton-X-100) for precisely 1 minute. Next an equal amount of 2 % (w/v) (1 % (w/v) final concentration) paraform aldehyde in PEM80 was added and incubated for 1 minute. The solution was removed and the cells were washed 3 times for 1 minute with PEM80 supplemented with 1 µM Paclitaxel (10-2095, Focus Biomolecules).

Finally, imaging buffer was applied to cavity slides. The imaging buffer contained ~2.5 - 5 nM labelled kinesin, 500 µM ATP, 1 mM DTT, 1 mM methyl viologen, 1 mM ascorbic acid, 20 µM Paclitaxel, 0.4 µg/µL Pyranose Oxidase (P4234-250UN, Thermo Fisher Scientific), ~96 U/mL Catalase (Bovine, 100429, MP Biomedical) in PEM80 supplemented with 10 % (w/v) glucose. The coverslip was then mounted with cells facing towards the cavity and sealed using Pico dent twinsil speed 22 (picodent Dental-Produktions-und Vertriebs GmbH) and curated at 37° C for 5 minutes.

### Live-cell kinesin tracking

U-2 OS cells were seeded at least two days prior to the experiments. The following day after seeding, 500 ng DNA of in Opti-MEM was mixed with 2 µL lipofectamine 2000 (11668019, Thermo Fisher Scientific) and incubated for 15 minutes. Afterwards, the mixture was applied dropwise to the cell culture. After 2 hours, the media was exchanged with fresh prewarmed FluoroBrite DMEM (A1896701, Thermo Fisher Scientific) and the cells were left over night to incubate.

On the experiment day, cells were labelled with 100 nM JF549 HaloTag ligand (HT1020, Promega) for 15 minutes in fresh prewarmed FluoroBrite. Then, the cells were washed 2 times for 15 minutes in FluoroBrite supplemented with 10 µM Paclitaxel. Afterwards, the coverslips were mounted and sealed as described previously.

## Data Analysis

The data analysis was designed by us and was implemented and executed in MATLAB R2021b. Some functions were inspired by ChatGPT code examples, in particular for the visualization of the data.

### DNA origami evaluation

Localization tracks with a photon number *N* > 3000 were split into tracks spanning a maximum photon number of *N*_max_ = 3000. Initial photon detections *N*_*c*_ used to reach the minimum FWHM of the respective E-PSFs were discarded. To ensure static single fluorophores at one binding site, the tracks were filtered by a range of the mean detection rate; a minimal number of detections per track; a maximum standard deviation in each dimension; and a maximum ellipticity in the lateral plane. The parameter histograms of the unfiltered and filtered tracks are shown in Supplementary Fig. S7. Supplementary Table T1 lists the corresponding parameter limits.

Due to separate lateral and axial updates in our 3D-MINSTED localization process, the center positions 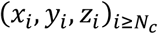 were split into sequences of *j* updates in the lateral plane (*x*_*j*_, *y*_*j*_) and along the axial direction (*z*_*j*_). The localization precisions were obtained by calculating the standard deviation 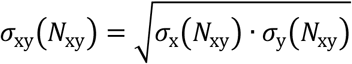 and σ_z_(*N*_z_) of the center positions averaged in blocks of *N*_*xy*_ photons for (*x*_*j*_, *y*_*j*_) and, respectively, *N*_z_ photons for (*z*_*j*_). For each valid localization precision estimate, we required at least 5 non-overlapping blocks. To ensure comparable results across the full range of photon numbers, tracks with no valid localization precision at *N*_min_ = 200 photons (equivalent of 5*N*_min_ = 1000 photons in both axial and lateral directions, respectively) were discarded. For the subsequent evaluation, the mean value of center positions 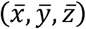 were used for clustering and fitting.

To estimate the binding sites, the retained center positions 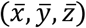 were grouped in clusters with the function *dbscan* (Matlab 2021b) maximally covering 8 nm in diameter in 3D. A secondary clustering was then performed with a maximum diameter of 34 nm to detect the 3×3 DNA origami grids. GATTAQuant specifies them with grid constant of 12 nm along one direction and 10.9 nm along the orthogonal direction. Clusters with less than six clustered binding sites and the corresponding tracks were discarded.

For each origami cluster, a 3×3 grid representing the expected binding sites, was matched to the measured cluster to get the cluster’s orientation in 3D and to estimates its grid constants. Next, the center positions were assigned to the nearest expected binding site, where center positions from the same track were allocated together to a cluster and binding site. Again, clusters with less than six binding sites and the corresponding tracks were discarded, resulting in a total of 87 matched and retained origamis. To overlay the matched clusters, they were recentred and rotated into a common orientation (*x*_rot_, *y*_rot_, *z*_rot_), where the larger grid constant was oriented along the x-axis and the smaller one along the y-axis. The retained clusters are shown in Supplementary Fig. S8.

For the retained tracks the localization precisions σ_xy_(*N*_xy_) and σ_z_(*N*_z_) were calculated as described above, based on the rotated data. Modelling those values with the function σ_est_(*N*) = *a*/(*b* + *N*)^*c*^ 0 < *a*, 0 < *b* and 0 < *c* < 0.5 estimates the lateral and axial localization precisions for the complete track^12^.

### Tracking evaluation

Tracks with at least 1000 detected photons and a standard deviation of at least 10 nm along the x- or y-axis were considered as tracks potentially stemming from moving kinesin. The initial detections before the E-PSFs reached their smallest values; detections during a further 20 ms interval; and detections during the final 20 ms were ignored to avoid intervals with insufficient localization precision and potential excursions due to background detections. The count rate was estimated as the reciprocal of the moving mean over time differences of consecutive photon detections with 100 binned values. Overly bright tracks with a mean count rate > 16000 cps (counts per second) were removed, because several fluorophores might have been tracked simultaneously. Tracks with a smaller count rate are probably either single fluorophore tracks or two fluorophore tracks with a bleaching event. To find a potential bleaching event from two to one fluorophore, an edge detection on the count rate was performed by using the function *ischange* (Matlab R2021b) with the ‘Statistic’ parameter set to ‘mean’ and the penalty parameter ‘MinThreshold’ = 1/3 · *n* · var{countrate}, where *n* denotes the number of countrate values in the track. If more than one bleaching step was found for a track, the track was removed, because it could stem from multiple fluorophores that were not removed by the previous mean count rate filter. Detections within the first 20 ms after the bleaching step were removed also to keep only single fluorophore tracks for the further analysis.

A principal component analysis was performed on the center positions of the retained fluorophore tracks and the coordinates were rotated along the principal axis in the lateral plane with a new coordinate system (*x*_*M*_, *y*_*M*_, *z*_*M*_), where the x_M_-axis is the direction of the principal axis. In order to remove static fluorophores, tracks briefer than *t*_min_ = 100 ms and tracks spanning less than 80 nm along the principal axis (i.e. less than 5 potential kinesin steps) were filtered out. Particularly for the live cell data, the trajectories along the microtubules were not linear probably due the flexibility and bending of the microtubules. Therefore, the shape of the microtubule was modelled with a third-degree polynomial along the orthogonal axes y_M_ and z_M_. The standard deviation of the distance from the microtubule model was calculated to obtain an initial estimate of the single-photon spatial precisions 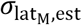 and 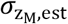 on the curved trajectory. Due to the rotational symmetry of the vortex donut, the single-photon spatial precisions along the principle-axis 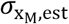 and orthogonal-axis 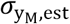 were assumed to be equal to 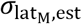. Tracks with large single-photon spatial precision σ _*i*, est_ > σ_max_ were filtered out. Such tracks may have been disturbed by other fluorophores moving through the sampling volume. The unfiltered and filtered tracks along with the corresponding filter parameters listed in Supplementary Table T2 and shown in Supplementary Fig. S9 and S12.

To identify kinesin steps, the retained tracks were segmented with a custom ‘pruned-exact-linear-time’ (PELT) algorithm^27,28^ on the center positions (*x*_*M,i*_ *y*_*M,i*_, *z*_*M,i*_), where the centers were normalized with the corresponding single-photon spatial precision in the corresponding axis. The PELT penalty parameter was set to 120 normalized spreads. Segments spanning less than 120 photons were flagged as unreliable due to large position uncertainty. Segments were also flagged as unreliable if the distance between the mean positions in segments was < 5 nm, probably caused by i) just the localization uncertainty, ii) several kinesin steps in rapid succession, leading to distances > 32 nm, or iii) if the step direction was not aligned with the principal axis. Finally, only tracks with at least three distance values were considered. The results for all segments and for sequential reliable segments only are shown in Supplementary Fig. S10 and S13 and the corresponding filter parameters are given in Supplementary Table T3.

The spatio-temporal resolution and the estimation of the SBR are introduced in reference^13^ and briefly described in the Supplementary information.

### Statistics and reproducibility

The measurements for the 3×3 DNA origami grids were performed for 7 different FOVs, resulting in a total of 599 localizations of the binding sites on 87 automatically detected grids in 6 different FOVs. Fig. 2b and 2c display grids of one exemplary FOV. Fig. 2h shows an overlay of all 87 detected grids with localizations based on 200 binned photons.

For the tracking data shown in Fig. 3 and 4, 7 fixated and 4 living cells with 20 and 25 FOVs were measured, resulting in 188 and 337 evaluated tracks with 788 and 1207 steps, respectively.

