## Supplementary Information for "3D-MINSTED nanoscopy and protein tracking in densely labelled living cells"

#### MINSTED microscope

*Main components*

| | |
|---|---|
| 488 | Diode laser 0488-06-01-0060-100 (60 mW cw), Cobolt, Solna, Sweden. |
| 532 | Pulsed fiber laser LDH-P-FA-530XL (> 200 mW, < 100 ps, pulsed picked), PicoQuant, Berlin, Germany. |
| 560 | Diode laser 561L-50-COL-PP (100 mW cw), Oxxius, Renens, Switzerland. |
| $636_{1,2,3}$ | Custom-built fiber amplified diode laser from laser diodes HL63391DG (633-643 nm, 200 mW), Ushio Opto Semiconductors, Tokyo, Japan; and a custom Praesodymium-doped fiber 220309/TB375-1 (Le Verre Fluoré, Bruz, France). |
| 855 | Super-luminescent LED module EBD273106-13 (< 5 mW cw), Exalos, Schlieren, Switzerland. |
| 1000 | Super-luminescent LED module SLD-1000-100-PM-25 (25 mW cw), Innolume, Dortmund, Germany. |
| $APD_{1,2,3}$ | Single-photon counting module SPCM-AQR-13, Excelitas, Wiesbaden, Germany. |
| PMT | Photonmultiplier DC-Modules MD963, PerkinElmer Optoelectronics, Shelton, USA. |
| $CAM_1$ | USB3 sCMOS camera pco.panda 4.2 (2048×2048 pixels 6.5×6.5 $\mu m^2$, 16 bits, 100 fps), PCO, Kelheim, Germany. |
| $CAM_2$ | USB3 CMOS camera xiQ MQ013RG-ON (1280×1024 pixels 4.8×4.8 $\mu m^2$, 8/10 bits, 210 fps), Ximea, Münster, Germany. |
| SLM | Spatial-light modulator X10468-01 (792×600 pixels 20×20 $\mu m^2$, 60 fps), Hamamatsu Photonics, Hamamatsu City, Japan; controller X10468-series (60 fps), Hamamatsu Photonics, Hamamatsu City, Japan. |
| $EOD_{x,y}$ | Electrooptic deflectors M-311A (ADP, Ø1.5 mm, 219 mm long, 3.5 µrad/V, 185 pF, ±500 V), Conoptics, Danbury, CT, USA; high-voltage Amplifier WMA-100 (±175 V, 100 mA, 500 kHz), Falco Systems, TH Katwijk aan Zee, Netherlands. |

| | |
|---|---|
| res. EOM | Resonant Electrooptic polarization modulator AM7R2-Vis_20 (20 MHz, Ø2 mm) QUBIG, München, Germany; wideband amplifier AMP590033H (2 W, 900 MHz) Becker, Asbach, Germany. |
| Flip mirror | Motorized Filter Flip Mount MFF101 (Ø1''), Thorlabs. |
| Deformable mirror | Hexagonally segmented deformable mirror DM HEX 111 (37 segments, 3.5 µm physical stroke, 7 µm wavefront stroke, Ø3.8 mm, 375 µm pitch, 100 kHz digital update rate) with controller X-CL140, Boston Micromachines Corporation, Cambridge, MA, USA. |
| Galvo scanner | Galvonometric mirrors 6 mm×10 mm, galvo 6215H, servo drivers 671, Cambridge Technology, Bedford, MA, USA. |
| $FPGA_{1,2}$ | PCIe board 7852R with drivers and software LabVIEW 2021, National Instruments, Austin, TX, USA. |
| Pulse Streamer | Synchronous digital pattern and arbitrary waveform generator Pulse Streamer 8/2 (digital output: 1 GS/s sampling rate; analog output: 125 MS/s, ±1 V voltage range, 14 bit), Swabian Instruments, Stuttgart, Germany. |
| Objective | 60×/1.3 NA silicon oil immersion objective UPLSAPO60XS2 Olympus, Tokyo, Japan. |
| Stages | Three-axis piezo stage P-733.3DD with controller E-712.3CD, Physik Instrumente, Karlsruhe, Germany; on three linear stages LNR502/M with controller BSC203, Thorlabs. |
| *Power control* | |
| $AOM_{1,2,3,4}$ | Acousto-optic modulator MT110-A1,5-VIS (110 MHz carrier drive frequency, Ø1.5 mm×2 mm) with RF driver MODA110, AA Sa, Orsay, France. |
| $LPC_{1,2}$ | Laser Power Controller LPC-VIS (Ø4 mm, 425 – 780 nm) with corresponding controller, Brockton Electro-Optics Corp., Bridgewater, MA, USA |
| $LPC_3$ | Laser Power Controller LPC-NIR (Ø4 mm, 700 – 1100nm) with corresponding controller, Brockton Electro-Optics Corp., Bridgewater, MA, USA |
| *Filters* | |
| $F_{1,2,3}$ | Edge basic long-pass filter 633 LP, Semrock, Rochester, NY, USA. |
| $F_4$ | BrightLine single-band band-pass filter FF01-582/75-25, Semrock. |
| $F_5$ | Edge transmission long-pass filter ET542lp, Chroma Technology, Bellows Falls, VT, USA. |
| $F_6$ | VersaChrome Edge tunable short-pass filter TSP01-628-25x36, Semrock. |
| $F_7$ | Razor-edge ultrasteep short-pass filter SP01-633RU, Semrock. |
| $F_8$ | Emission filter long-pass HQ580lp, Chroma Technology, Bellows Falls, VT, USA. |
| $F_9$ | Emission filter band-pass HQ580/40M, Chroma Technology. |
| $F_{10}$ | Standard filter band-pass BP610-105, Ferroperm Optics A/S, Vedbæk, Denmark. |
| $F_{11}$ | Notch filter ZET633NF, Chroma Technology. |
| $F_{12}$ | Raman emitter long-pass RET493lp, Chroma Technology. |
| $F_{13}$ | Emission filter band-pass HQ500/40M, Chroma Technology. |
| $F_{14}$ | Emission filter band-pass HQ510/20M, Chroma Technology. |
| $F_{15}$ | Hard-coated band-pass filter FBH850-40, Thorlabs. |
| ID | Iris diaphragms D25S, Thorlabs. |
| Pin. | Motorized pinhole wheel MPH16-A, Thorlabs. |
| VPP | Vortex phase plate V-633-10-1, Vortex Photonics, München, Germany. |

*Lenses*

| | |
|---|---|
| $C_{1,2}$ | Fiber collimator 15 mm, 60FC-A15-4-01, Schäfter+Kirchhoff, Hamburg, Germany. |
| $C_{3,4}$ | Fiber collimator 4.5 mm, 60FC-A4,5-4-01, Schäfter+Kirchhoff. |
| $C_{7,8,9,14,15,16}$ | Fiber collimator 7.5 mm, 60FC-A7,5-4-01, Schäfter+Kirchhoff. |
| $C_{5,6,12,13}$ | Fiber collimator 7.5 mm, 60FC-A7.5-4-02, Schäfter+Kirchhoff. |
| $C_{10,11}$ | Fiber collimator 18 mm, 60FC-A18-4-01, Schäfter+Kirchhoff. |
| $L_1$ | Achromatic doublets 300 mm, 322273322, Linos Photonics, Göttingen, Germany. |
| $L_{2,3,4}$ | Achromatic doublet 100 mm, 49-333, Edmund Optics, Mainz, Germany. |
| $L_5$ | Achromatic doublets 80 mm, AC254-080-A, Thorlabs. |
| $L_6$ | Achromatic doublets 250 mm, 322272322, Linos Photonics. |
| $L_7$ | Achromatic doublets 140 mm, 322351000, Linos Photonics. |
| $L_{8,9}$ | Achromatic doublets 300 mm, AC254-300-A, Thorlabs. |
| $L_{10}$ | Achromatic doublets 60 mm, 312332000, Linos Photonics. |
| $L_{11}$ | Achromatic doublets 100 mm, 312333000, Linos Photonics. |
| $L_{12}$ | Achromatic doublets 200 mm, 312405000, Linos Photonics. |
| $L_{13}$ | Achromatic doublets 300 mm, 322273322, Linos Photonics. |
| $L_{14}$ | Achromatic doublets 150 mm, AC254-150-A, Thorlabs. |
| $L_{15,16}$ | Achromatic doublets 200 mm, AC254-200-A, Thorlabs. |
| $L_{17,18}$ | Achromatic doublets 60 mm, 312316000, Linos Photonics. |
| $L_{19}$ | Achromatic doublets 250 mm, 322272322, Linos Photonics. |
| $L_{20,21}$ | Achromatic doublets 200 mm, NIR, 322367525, Linos Photonics. |

*Mirrors*

| | |
|---|---|
| CM | Cold mirror M254C00, Thorlabs. |
| Pel | Removable pellicle beamsplitter BP145B1 (R/T ratio 45/55), Thorlabs. |
| $DM_1$ | Dichroic mirror ZT594RDC, Chroma Technology. |
| $DM_{2,3}$ | Dichroic mirror TFPB-532+637-45+HRs450-750/45/ARPW2037UV, Laser Components Germany GmbH, Olching, Germany. |
| $DM_4$ | Multi band dichroic mirror ZET405/488/561/640RPC, Chroma Technology. |
| $DM_5$ | Dichroic mirror ZT532RDC, Chroma Technology. |
| $DM_6$ | Dichroic mirror FF509-FDi01-25x36, Semrock. |
| $DM_7$ | Dichroic mirror DMLP900, Thorlabs. |
| Parabolic mirror | Off-axis parabolic mirror, MPD120-P01, Thorlabs. |
| Unmarked | MaxMirror ultra-broadband mirror, Semrock. |

*Polarization optics*

| | |
|---|---|
| $PBS_{1,2,3,4,5,6,7}$ | Polarizing beam splitter cube PTW 0.20, Bernhard Halle, Berlin, Germany. |
| $PBS_{8,10}$ | Polarizing beam splitter cube PTW 1.20, Bernhard Halle. |
| $PBS_{9,11}$ | Polarizing beam splitter cube PTW 2.20, Bernhard Halle. |
| $\lambda/4$ | Achromatic quarter-wave retarder plate RAC 3.4.10, Bernhard Halle. |
| $\lambda/2$ | Achromatic half-wave retarder plate RAC 3.2.10, Bernhard Halle. |
| $\lambda/2_s$ | Segmented achromatic half-wave retarder plate RAC 3.2.10, Bernhard Halle. |
| $W_1$ | Wollaston prism 68-820, Edmund Optics. |

*Fibers*

| | |
|---|---|
| $PM_{1,2,3}$ | Polarizing maintaining fiber P3-630PM-FC5, Thorlabs. |
| $PM_4$ | Polarizing maintaining fiber P3-488PM-FC5, Thorlabs. |
| $PM_{5,6}$ | Polarizing maintaining fiber P3-488PM-FC10, Thorlabs. |
| $PM_7$ | Polarizing maintaining fiber P3-980PM-FC5, Thorlabs. |
| $GI_1$ | Graded-index multimode fiber M115L01 (Ø50 µm), Thorlabs. |
| $GI_{2,3}$ | Graded-index multimode fiber M31L01 (Ø62.5 µm), Thorlabs. |

### Calibration of the defocus shift

To calibrate the positional shift along the axial direction, a sample of dried Poly-L-Lysin (P8920-100ML, Sigma-Aldrich) with 100 nm gold beads (EM.GC100/7, BBI Solutions) on the cover slip, sealed in Mowiol (81381-250G, Sigma Aldrich) was prepared. In a 1.5 $\mu$m $\times$ 1.5 $\mu$m region a single bead was scanned five times via the deformable mirror in the xz-plane with a STED laser beam from either STED Z1 or STED Z2. The wavefront was modulated by the SLM with a top-hat phase mask to create a 3D donut in the focal plane. The minimum of the donut was approximately centered in the region of interest and was used as a reference point. The scattered light was detected by the photon multiplier tube. To average out fluctuations, each image stack, containing five images, was summed up. To extract the minimum, the profiles ±30 nm around the central line of the x-axis were extracted and summed along the x-direction. In order to find the shift of the minimum along the axial direction, the minimum of the profile was fitted to a parabola.
Afterwards, the scan was repeated with a modulated wavefront to shift the minimum along the axial direction using the defocus Zernike polynomial[1] with four other different coefficients covering a distance of $\approx$ 100 nm from the focal plane. To calibrate the defocus shift scale, the measured values were linearly regressed. Supplementary Fig. S5 shows the defocus coefficients with the corresponding shifts and the respective linear fit.

### Determination of the spatiotemporal resolution

The spatial resolution was calculated by the standard deviation of the segmented mean position of the filtered tracks in the respective direction (Methods). To estimate the temporal resolution, positions along the principal axis $x_M$ within 20 ms of an estimated step were scaled, such that the pre-step mean position was zero and the post-step mean position was one. Mean positions larger than 20% deviation from the corresponding plateau were neglected to avoid artifacts due to insufficient data quality. To fine tune the instant of the kinesin step, the scaled steps were modelled with the function

$$x_M(t) = \max\left\{0; 1 - \exp\left(-\frac{t-t_0}{\tau}\right)\right\},$$

where $t_0$ is the instant the kinesin stepped and $\tau$ is the response time of the 3D-MINSTED localization. Steps with estimated $\hat{\tau} > 9$ ms were discarded for this analysis, because overly slow responses were probably due to off-axis steps. The scaled positions of the retained steps were translated by the estimated instants $t_0$ to the common time interval $[-T, T]$ with $T = 20$ ms and were overlayed. These aligned steps were then averaged with a spacing of $T/500$ and the average step was fit with the function $1 - \exp\left(-\frac{t}{\tau}\right)$ for $t \geq 0$ to get the final estimation of the response time $\tau$. A more detailed explanation is given in reference[2]. The spatial and temporal resolution data is shown in Supplementary Fig. S14.

### Signal-to-background ratio estimation

The signal-to-background ratio (SBR) was estimated with the tracks of the corresponding field of view (FOV). Detections at the end of a track $[T_{\mathrm{ter}} - t_{\mathrm{off}}, T_{\mathrm{ter}}]$, where $T_{\mathrm{ter}}$ is the termination time, with a time difference $\Delta t_i > t_{\mathrm{off}}/N_{\mathrm{off}}$ between two consecutive detections were considered as background photons $N_{\mathrm{bgr}}$. The background rate for all evaluated tracks in a FOV was defined as $k_{\mathrm{bgr}} = \langle N_{\mathrm{bgr},j}/t_{\mathrm{off},j} \rangle_{j \in \mathrm{FOV}}$. For the overall detection rate $k$, the number of detected photons of an evaluated track were divided by the total time of the localization. The SBR was computed as $(k - k_{\mathrm{bgr}})/k_{\mathrm{bgr}}$ as defined in reference[2]. All SBR histograms are shown in Supplementary Fig. S15.

### Supplementary tables and figures

#### Filter parameters

Supplementary Table T1 | Parameters for filtering the raw tracks.

| Sample | Number of photons | Detection rate [kcps] | | Standard deviation | | | Ellipticity | |
|---|---|---|---|---|---|---|---|---|
| | | min | max | $\sigma_x$ [nm] | $\sigma_y$ [nm] | $\sigma_z$ [nm] | min | max |
| 3×3 12 nm grid | ≥ 250 | ≥ 7 | ≤ 18 | ≤ 7 | ≤ 7 | ≤ 14 | $0.8\ \sigma_x \leq \sigma_y$ | $\sigma_y \leq 1.25\ \sigma_x$ |
| tracking in fixed cells | ≥ 1000 | - | - | ≥ 10 | ≥ 10 | - | - | - |
| tracking in live cells | ≥ 1000 | - | - | ≥ 10 | ≥ 10 | - | - | - |

Supplementary Table T2 | Parameters for filtering tracks.

| Sample | Distance along principal axis [nm] | Duration [ms] | Estimated standard deviation | | Average kinesin speed [nm/s] |
|---|---|---|---|---|---|
| | | | $\sigma_{xy,\mathrm{est}}$ [nm] | $\sigma_{z,\mathrm{est}}$ [nm] | |
| tracking in fixed cells | ≥ 80 | ≥ 100 | ≤ 10 | ≤ 25 | ≤ 1000 |
| tracking in live cells | ≥ 80 | ≥ 100 | ≤ 10 | ≤ 25 | ≤ 1000 |

Supplementary Table T3 | Filter parameters for step detection.

| Sample | Photon number per plateau | Direction along principal axis | Distance between plateaus | |
|---|---|---|---|---|
| | | | $d_{\mathrm{min}}$ [nm] | $d_{\mathrm{max}}$ [nm] |
| steps of kinesin in fixed/live cells | ≥ 120 | $\bar{x}_{M,\mathrm{center}_\mathrm{i}} \leq \bar{x}_{M,\mathrm{center}_{i+1}}$ | ≥ 5 | ≤ 32 |

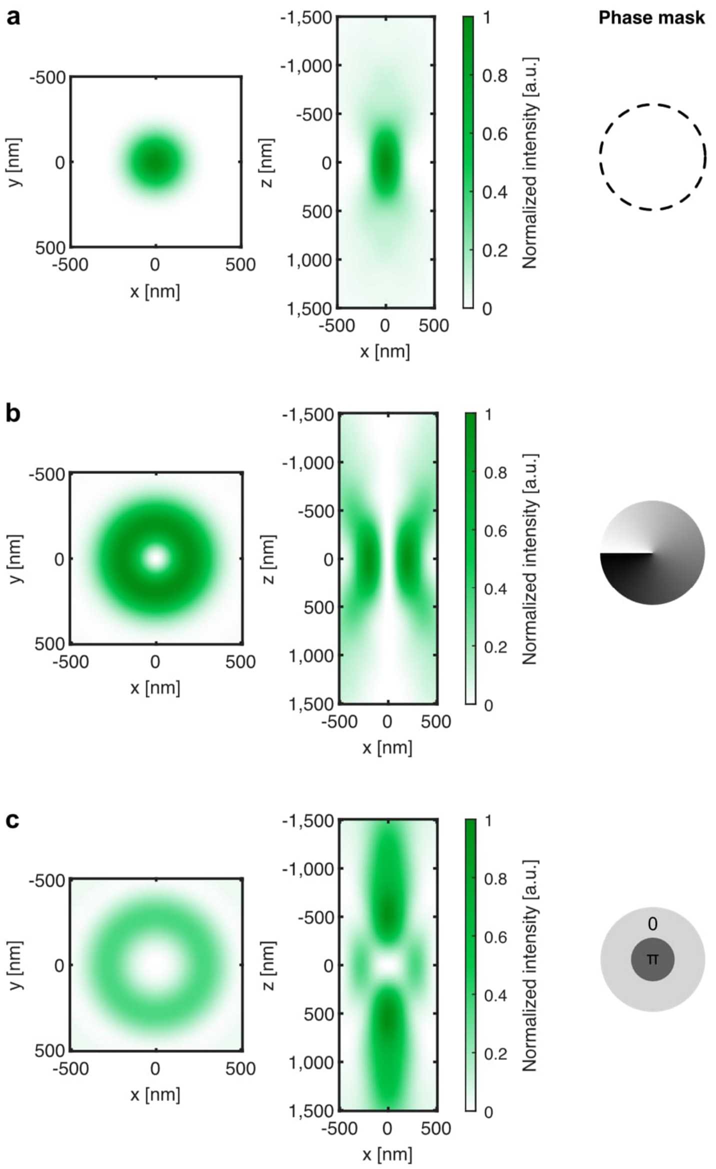


Suppl. Fig. S1 | Calculated excitation point spread functions. Calculated excitation PSFs[3] of an excitation beam with a wavelength of 580 nm for a Gaussian beam and 2D/3D donut in the lateral and axial plane with the corresponding phase mask.

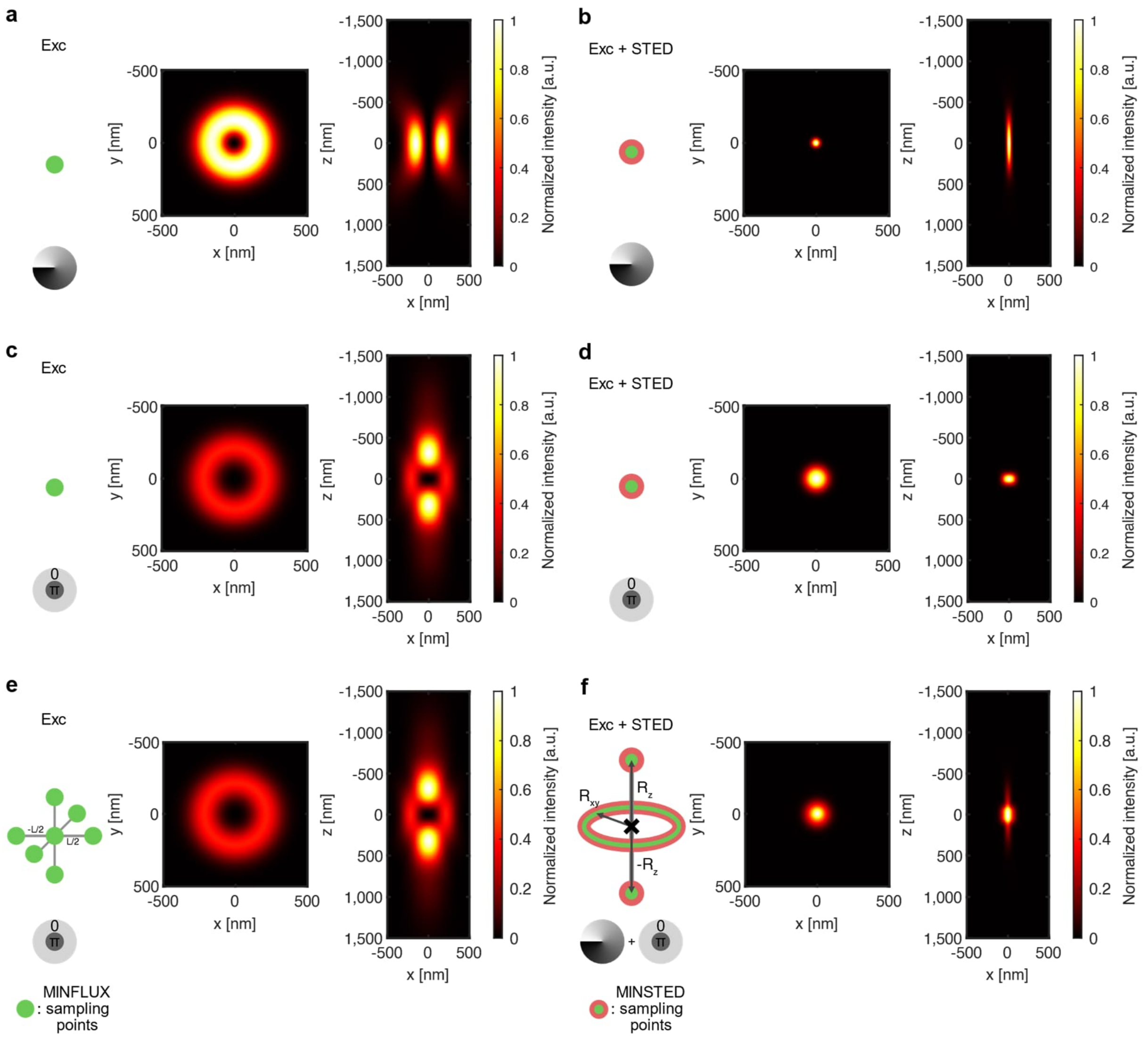


Suppl. Fig. S2 | Excitation volume from calculated point spread functions for the parameters of the fluorophore Cy3B for a single sampling position and for a measurement sampling pattern in MINFLUX and MINSTED. Calculated E-PSFs[3] of a single sampling position for a MINFLUX measurement within our setup boundaries at an excitation wavelength of 580 nm and a power of 100 µW with a 2D-donut (a), generated by a vortex phase plate, and with a 3D-donut (c), generated by a tophat phase mask. For a MINSTED measurement, the fluorophore is excited by a Gaussian beam with a wavelength of 532 nm, overlapped with 2D (b) and 3D (d) STED donuts with a pulse energy of $E_{xy}$ = 0.6 nJ and $E_z$ = 0.4 nJ at a wavelength of 636 nm. In (e) the effective excitable volume of a typical sampling pattern with a 3D-excitation donut from the commercial MINFLUX (Abberior Instruments) with a L=40 nm is calculated. Correspondingly, a typical sampling pattern of our MINSTED setup (f), described in 3D-MINSTED principle, are shown with $R_{xy}$=18 nm and $R_z$=40 nm and the resulting calculated point spread functions.

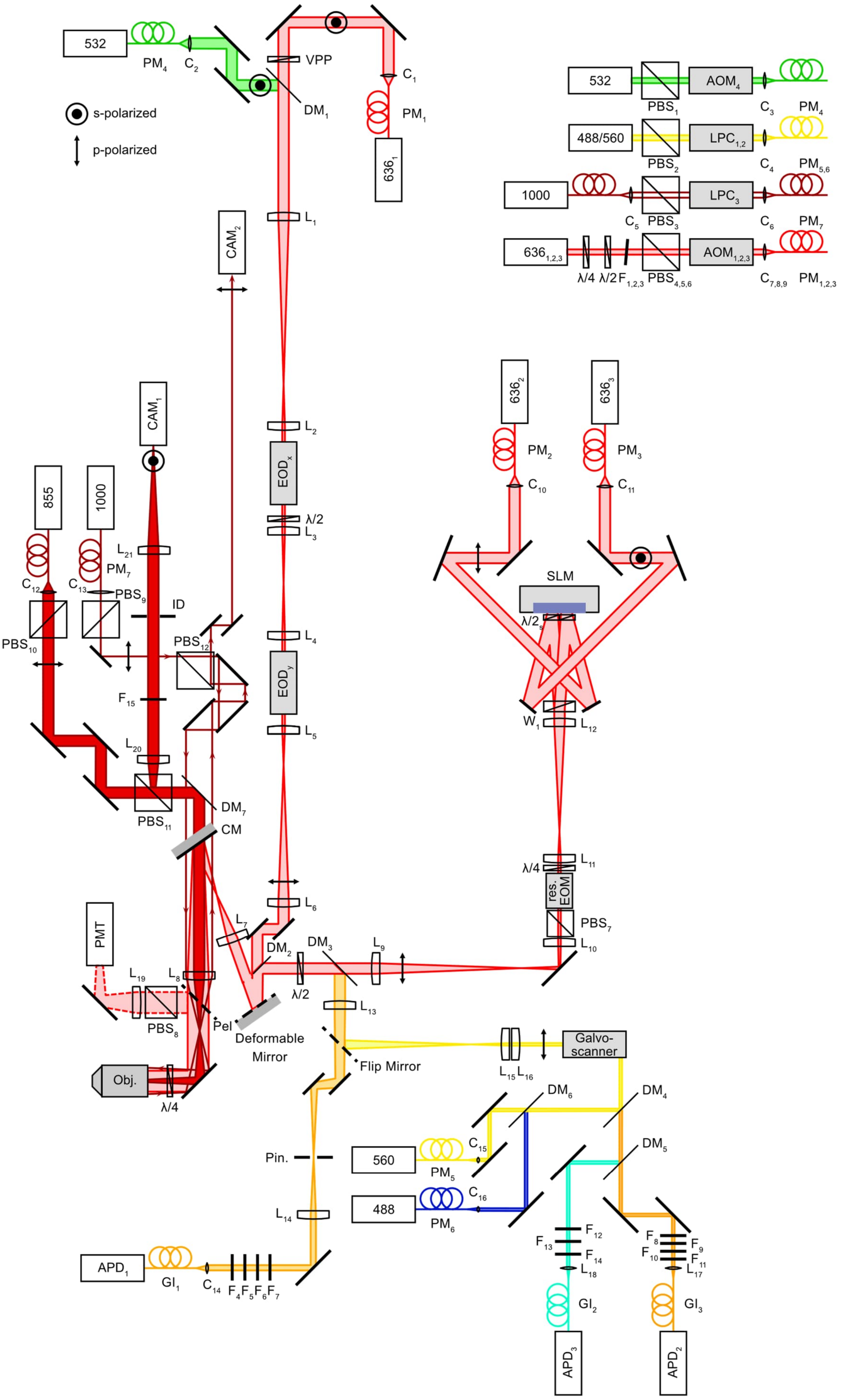
532
PM4
C2
VPP
DM1
C1
PM1
6361
s-polarized
p-polarized
532
PBS1
AOM4
C3
PM4
488/560
PBS2
LPC1,2
C4
PM5,6
1000
C5
PBS3
LPC3
C6
PM7
6361,2,3
λ/4
λ/2
F1,2,3
PBS4,5,6
AOM1,2,3
C7,8,9
PM1,2,3
L1
CAM2
L2
EODx
λ/2
L3
L4
EODy
L5
6362
6363
PM2
PM3
C10
C11
SLM
λ/2s
W1
L12
CAM1
855
1000
L21
PM7
C12
C13
PBS9
ID
PBS10
PBS12
F15
L20
PBS11
DM7
CM
L11
λ/4
res. EOM
PBS7
L10
L6
L7
DM2
DM3
L9
λ/2
PMT
L19
L8
PBS8
Pel
Deformable Mirror
L13
Flip Mirror
L15
L16
Galvo-scanner
Obj.
λ/4
DM6
DM4
560
C15
PM5
DM5
Pin.
488
C16
PM6
L14
F12
F13
F14
L18
F8
F9
F10
F11
L17
APD1
GI1
C14
F4F5F6F7
GI2
GI3
APD3
APD2

Suppl. Fig S3 | Outline of the MINSTED microscope. The MINSTED setup for localizations in 3D. The flip mirror can be flipped into the beam path for a confocal microscopy modality. The pellicle can be placed manually into the beam path to partially reflect the detected light onto the photon multiplier tube. The power modulations of the corresponding lasers are sketched in the upper right corner.

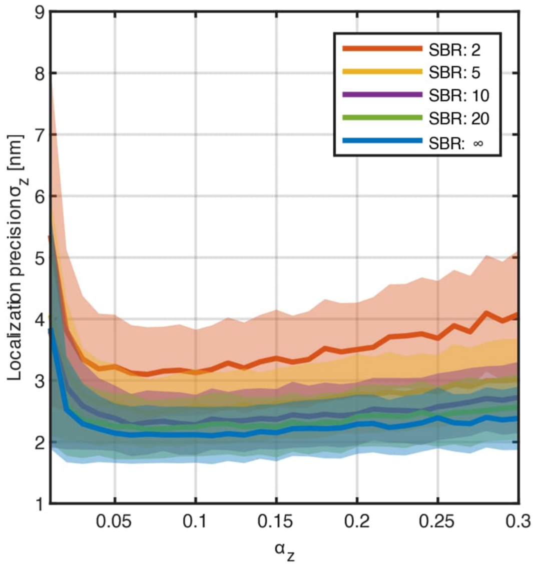


Suppl. Fig S4 | Localization precision $\boldsymbol{\sigma_z}$ vs. $\boldsymbol{\alpha_z}$ on simulated tracks. Localization precision $\sigma_z$ for 200 binned photons along the axial direction depending on the update step $\alpha_z$ for different SBR (line: mean of blocks with 200 photons on 1000 simulated tracks (Methods); shaded areas: ±1 standard deviations).

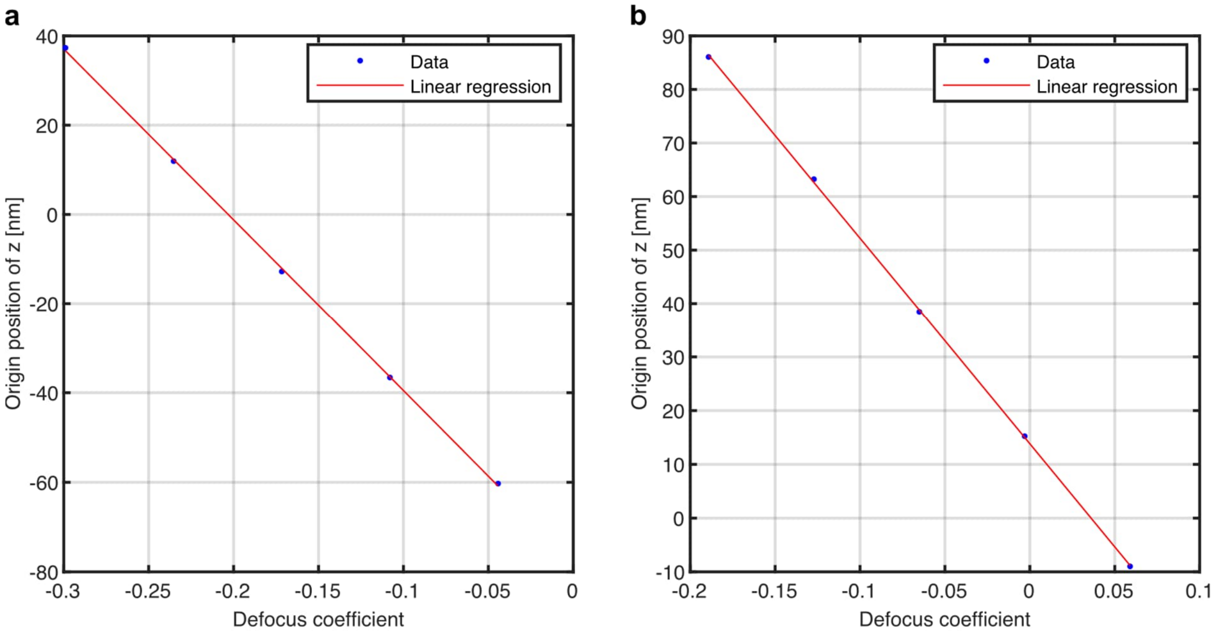


Suppl. Fig S5 | Calibration of the defocus coefficient with the shift along the axial direction. Position of the minimum of the STED beam depending on the defocus coefficient for STED Z1 (a) and STED Z2 (b).

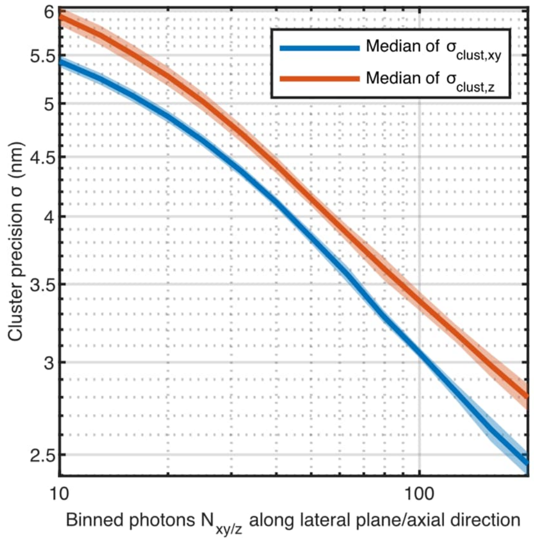


Suppl. Fig. S6 | Cluster precision versus number of binned photons in the lateral plane and axial direcion.

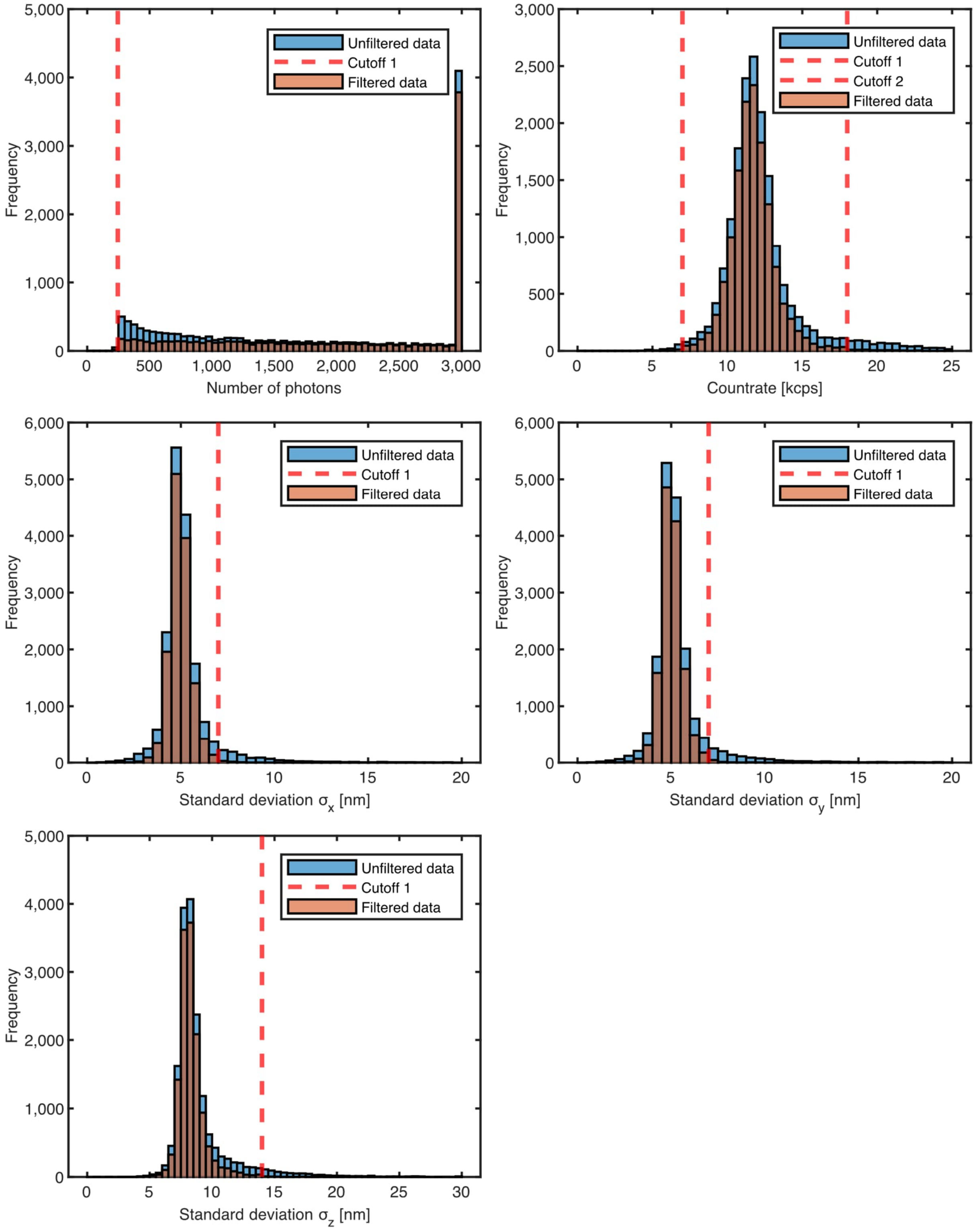


Suppl. Fig. S7 | Characteristic for tracks of 3×3 12 nm DNA origami grid. Histograms of unfiltered and filtered tracks for the filter parameters listed in Supplementary Table T1.

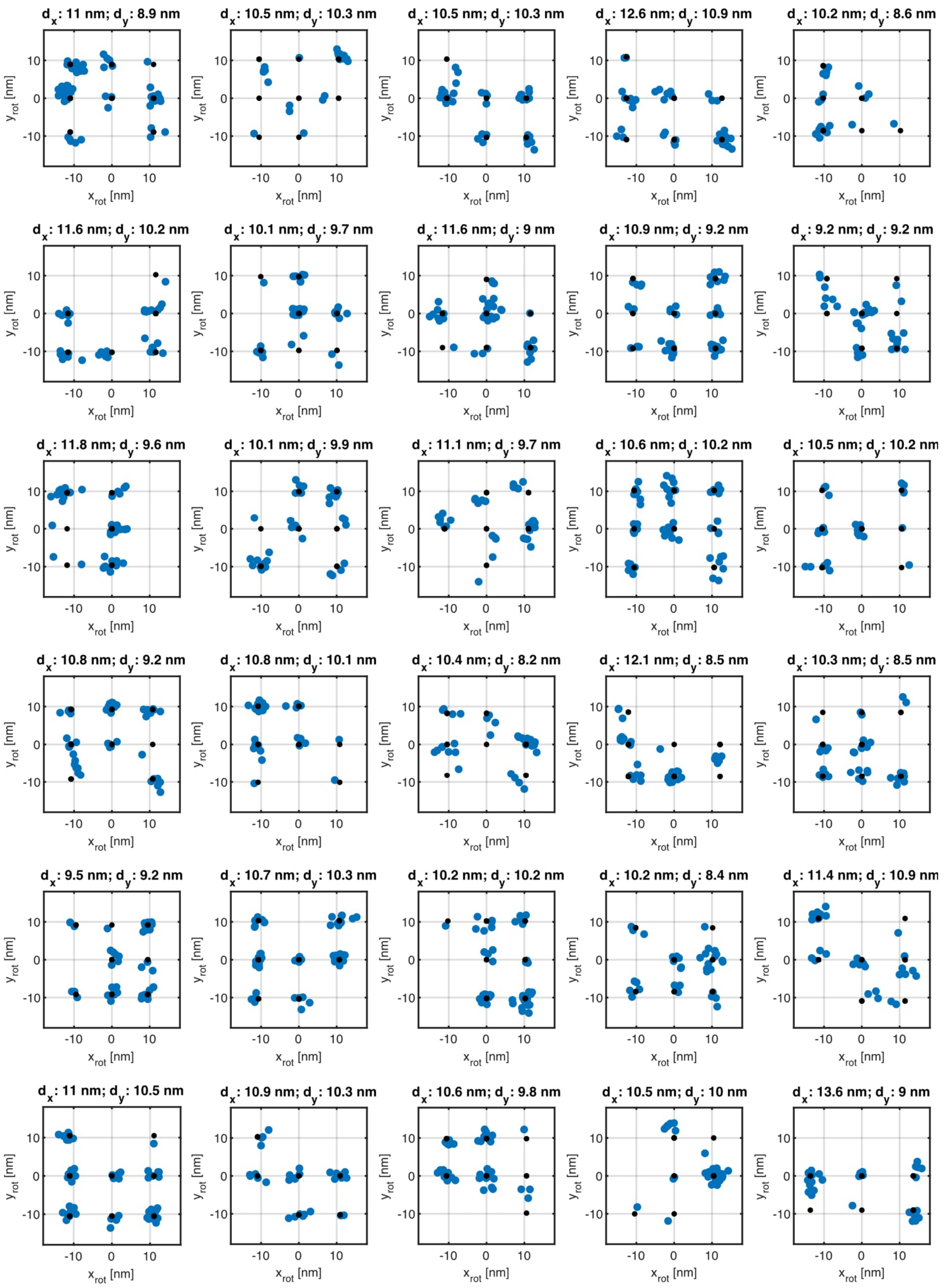

$d_x$: 11 nm; $d_y$: 8.9 nm
$d_x$: 10.5 nm; $d_y$: 10.3 nm
$d_x$: 10.5 nm; $d_y$: 10.3 nm
$d_x$: 12.6 nm; $d_y$: 10.9 nm
$d_x$: 10.2 nm; $d_y$: 8.6 nm
$d_x$: 11.6 nm; $d_y$: 10.2 nm
$d_x$: 10.1 nm; $d_y$: 9.7 nm
$d_x$: 11.6 nm; $d_y$: 9 nm
$d_x$: 10.9 nm; $d_y$: 9.2 nm
$d_x$: 9.2 nm; $d_y$: 9.2 nm
$d_x$: 11.8 nm; $d_y$: 9.6 nm
$d_x$: 10.1 nm; $d_y$: 9.9 nm
$d_x$: 11.1 nm; $d_y$: 9.7 nm
$d_x$: 10.6 nm; $d_y$: 10.2 nm
$d_x$: 10.5 nm; $d_y$: 10.2 nm
$d_x$: 10.8 nm; $d_y$: 9.2 nm
$d_x$: 10.8 nm; $d_y$: 10.1 nm
$d_x$: 10.4 nm; $d_y$: 8.2 nm
$d_x$: 12.1 nm; $d_y$: 8.5 nm
$d_x$: 10.3 nm; $d_y$: 8.5 nm
$d_x$: 9.5 nm; $d_y$: 9.2 nm
$d_x$: 10.7 nm; $d_y$: 10.3 nm
$d_x$: 10.2 nm; $d_y$: 10.2 nm
$d_x$: 10.2 nm; $d_y$: 8.4 nm
$d_x$: 11.4 nm; $d_y$: 10.9 nm
$d_x$: 11 nm; $d_y$: 10.5 nm
$d_x$: 10.9 nm; $d_y$: 10.3 nm
$d_x$: 10.6 nm; $d_y$: 9.8 nm
$d_x$: 10.5 nm; $d_y$: 10 nm
$d_x$: 13.6 nm; $d_y$: 9 nm
$x_{rot}$ [nm]
$y_{rot}$ [nm]

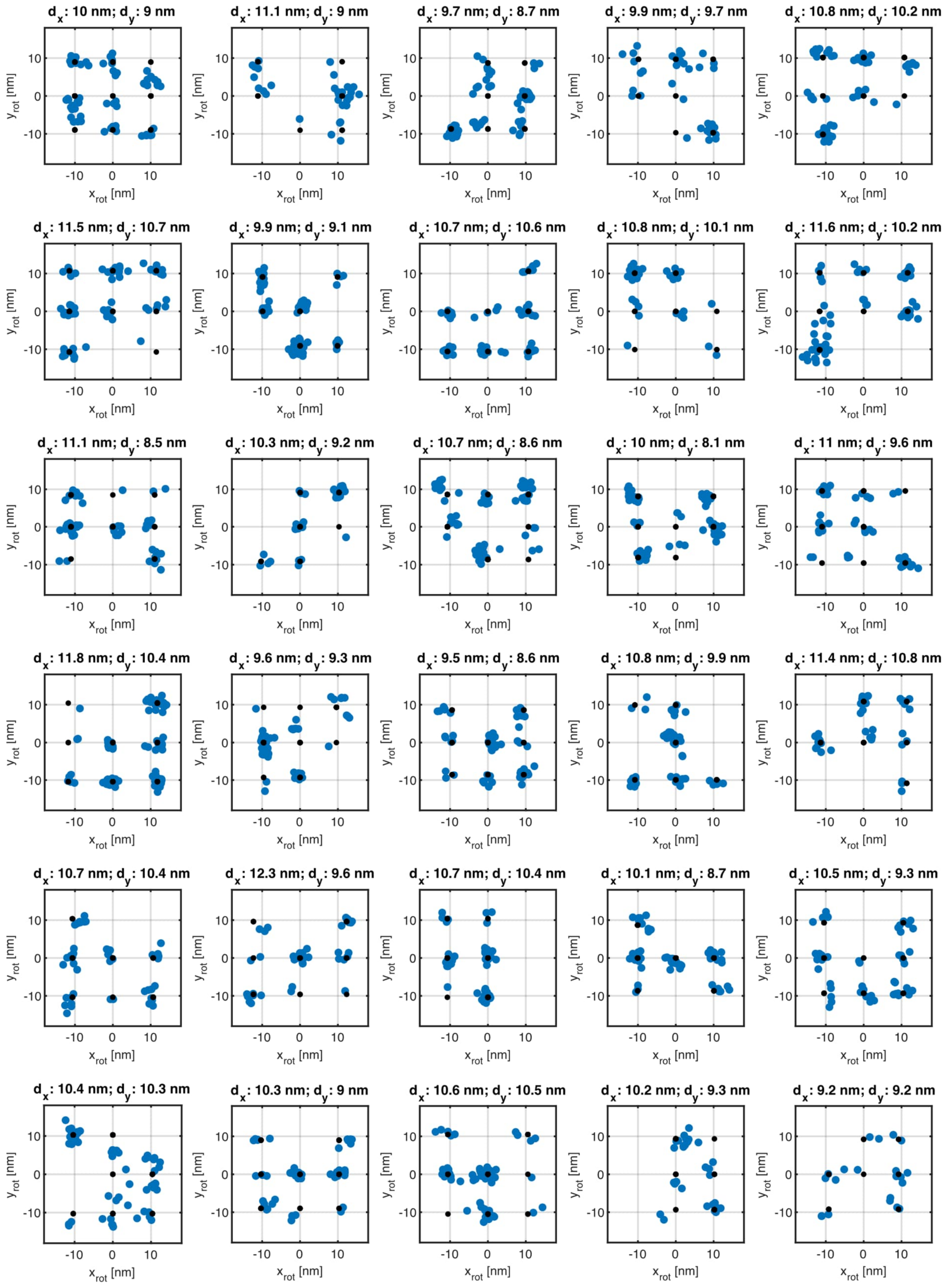
$d_x$: 10 nm; $d_y$: 9 nm
$d_x$: 11.1 nm; $d_y$: 9 nm
$d_x$: 9.7 nm; $d_y$: 8.7 nm
$d_x$: 9.9 nm; $d_y$: 9.7 nm
$d_x$: 10.8 nm; $d_y$: 10.2 nm
$d_x$: 11.5 nm; $d_y$: 10.7 nm
$d_x$: 9.9 nm; $d_y$: 9.1 nm
$d_x$: 10.7 nm; $d_y$: 10.6 nm
$d_x$: 10.8 nm; $d_y$: 10.1 nm
$d_x$: 11.6 nm; $d_y$: 10.2 nm
$d_x$: 11.1 nm; $d_y$: 8.5 nm
$d_x$: 10.3 nm; $d_y$: 9.2 nm
$d_x$: 10.7 nm; $d_y$: 8.6 nm
$d_x$: 10 nm; $d_y$: 8.1 nm
$d_x$: 11 nm; $d_y$: 9.6 nm
$d_x$: 11.8 nm; $d_y$: 10.4 nm
$d_x$: 9.6 nm; $d_y$: 9.3 nm
$d_x$: 9.5 nm; $d_y$: 8.6 nm
$d_x$: 10.8 nm; $d_y$: 9.9 nm
$d_x$: 11.4 nm; $d_y$: 10.8 nm
$d_x$: 10.7 nm; $d_y$: 10.4 nm
$d_x$: 12.3 nm; $d_y$: 9.6 nm
$d_x$: 10.7 nm; $d_y$: 10.4 nm
$d_x$: 10.1 nm; $d_y$: 8.7 nm
$d_x$: 10.5 nm; $d_y$: 9.3 nm
$d_x$: 10.4 nm; $d_y$: 10.3 nm
$d_x$: 10.3 nm; $d_y$: 9 nm
$d_x$: 10.6 nm; $d_y$: 10.5 nm
$d_x$: 10.2 nm; $d_y$: 9.3 nm
$d_x$: 9.2 nm; $d_y$: 9.2 nm
$x_{rot}$ [nm]
$y_{rot}$ [nm]

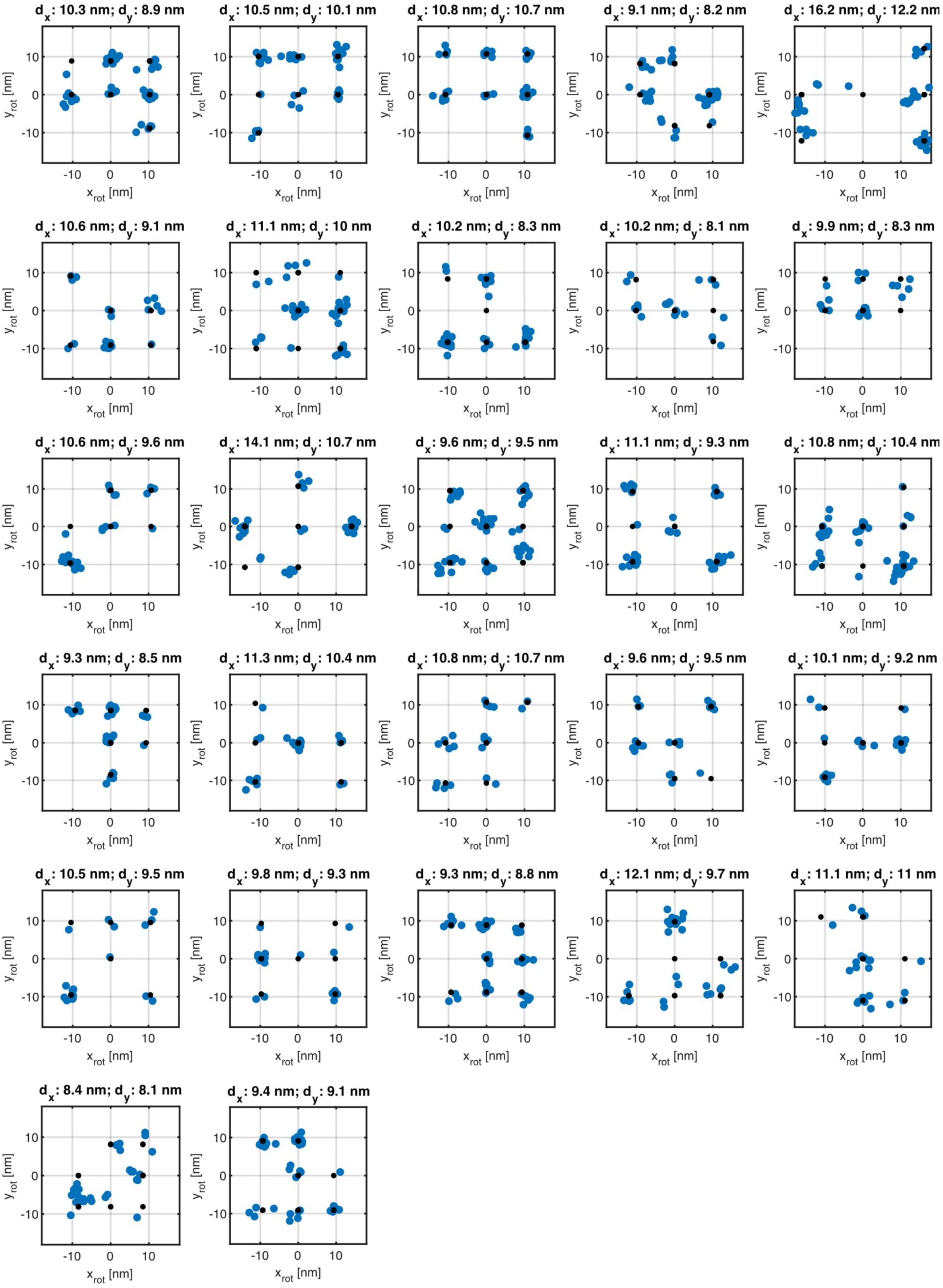


Suppl. Fig. S8 | Rotated DNA origami grids detected algorithmically. All 87 automatically detected DNA origami grids rotated into the common orientation ($x_{rot}$, $y_{rot}$, $z_{rot}$) with the corresponding fitted grid constants $d_x$ and $d_y$ (Methods).

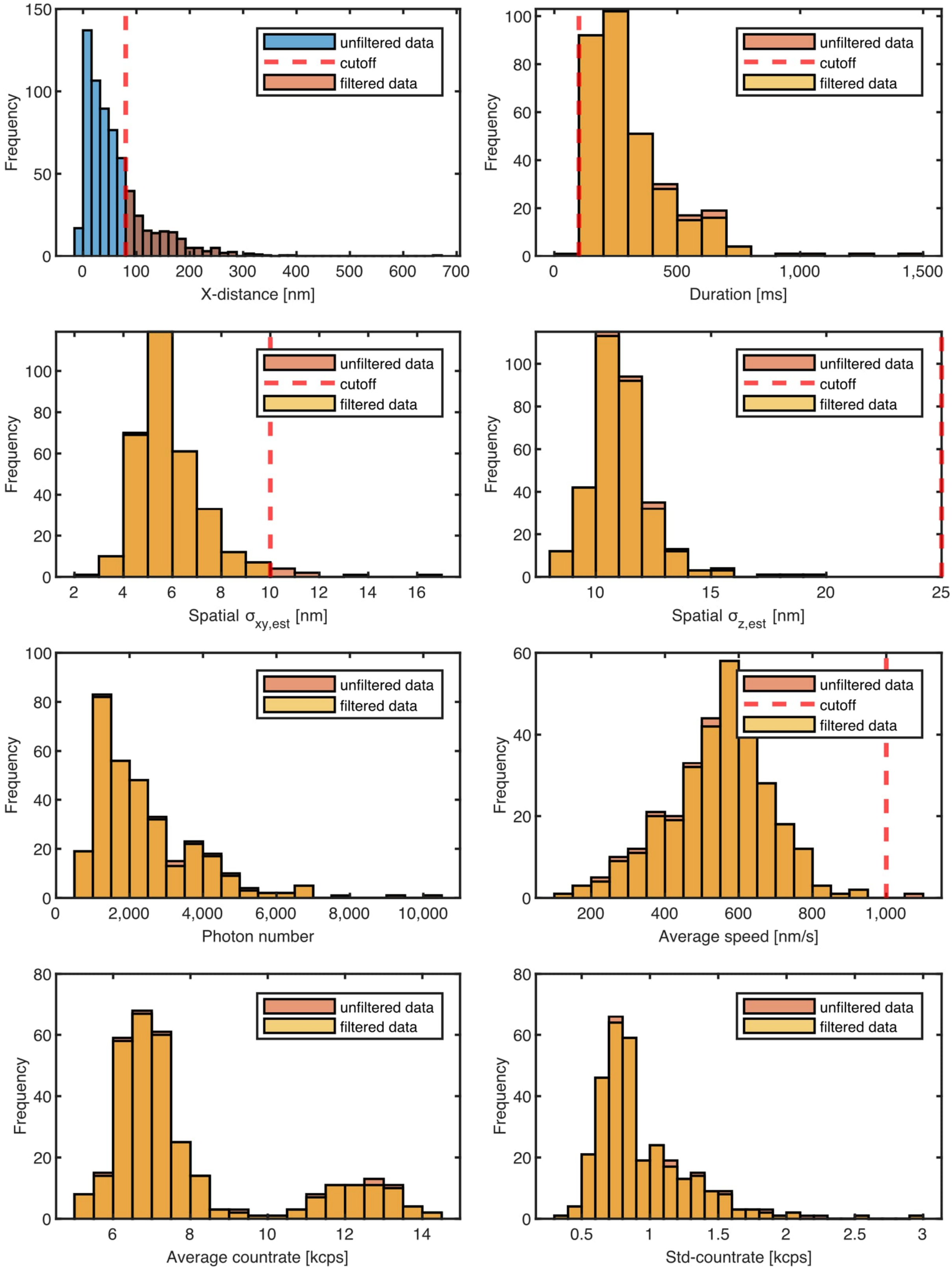


Suppl. Fig. S9 | Characteristic of fluorescence tracks measured by tracking of kinesin in fixed cells. Histograms of unfiltered and filtered tracks of kinesin tracking in fixed cells with filter parameters listed in Supplementary Table T2. First, the tracks were filtered by the distance along the principal axis (x-axis) and the retained tracks were filtered by the other filter parameters.

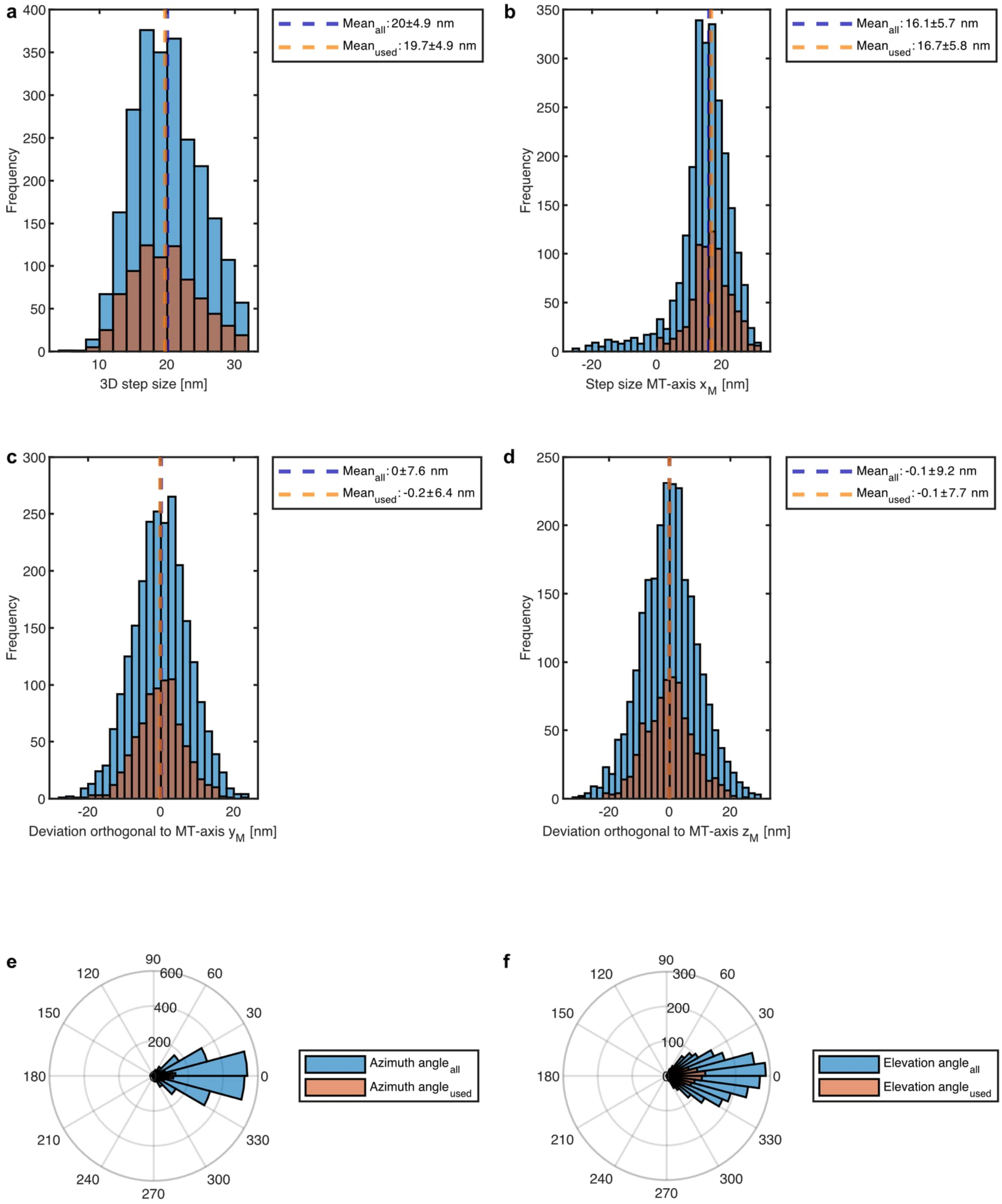


Suppl. Fig. S10 | Characteristic of detected steps in kinesin tracks of fixed cells. Histograms of all steps and of retained steps in kinesin tracks established in fixed along with their corresponding mean value values. The filter parameters are listed in Supplementary Table T3. (a) Euclidean step size distribution in 3D. (b) Step size along the principal axis and (c,d) the deviation along the orthogonal axis. (e,f) Distribution of azimuth and elevation angle.

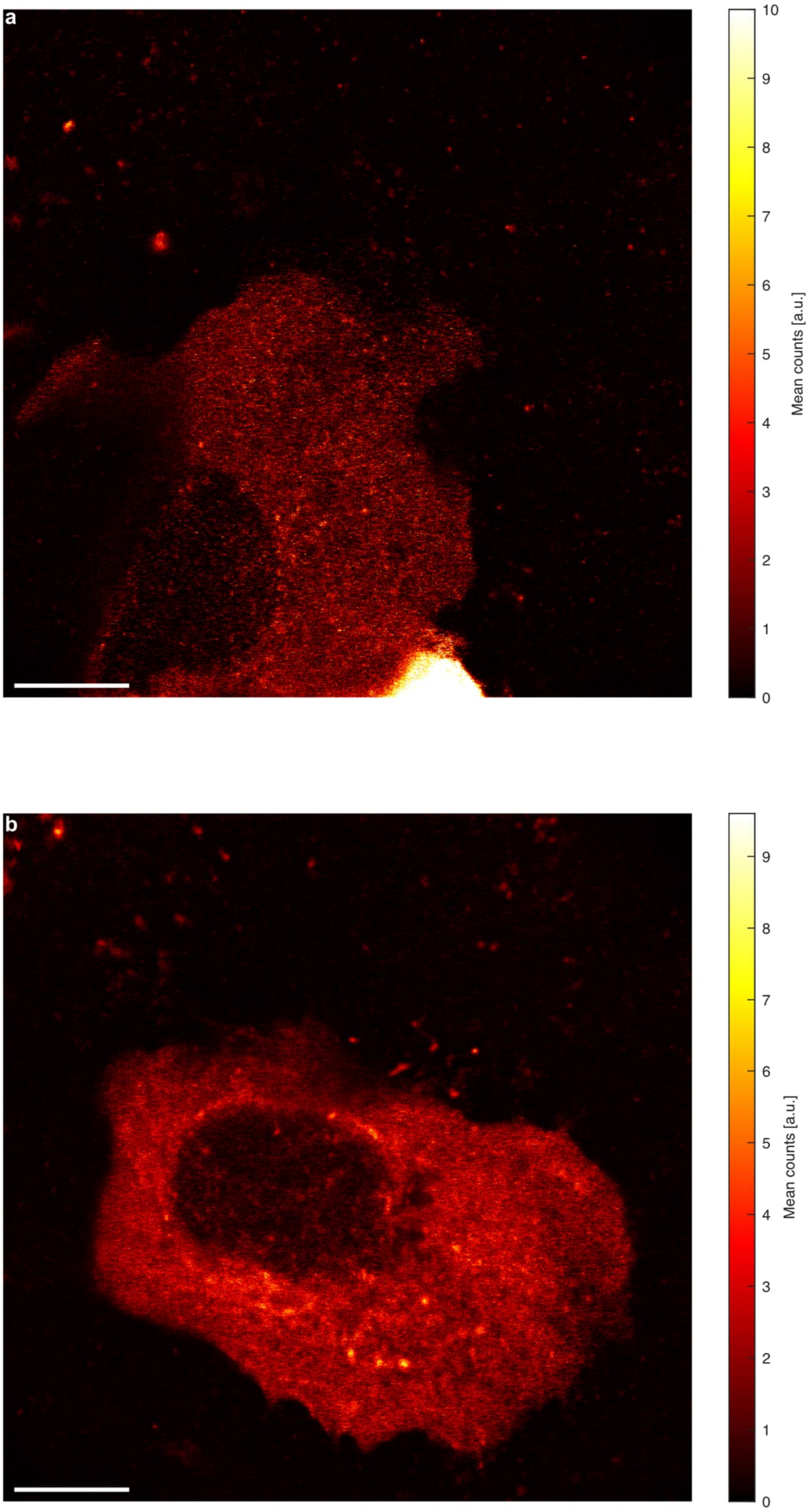
a
10
9
8
7
6
5
4
3
2
1
0
Mean counts [a.u.]
b
9
8
7
6
5
4
3
2
1
0
Mean counts [a.u.]

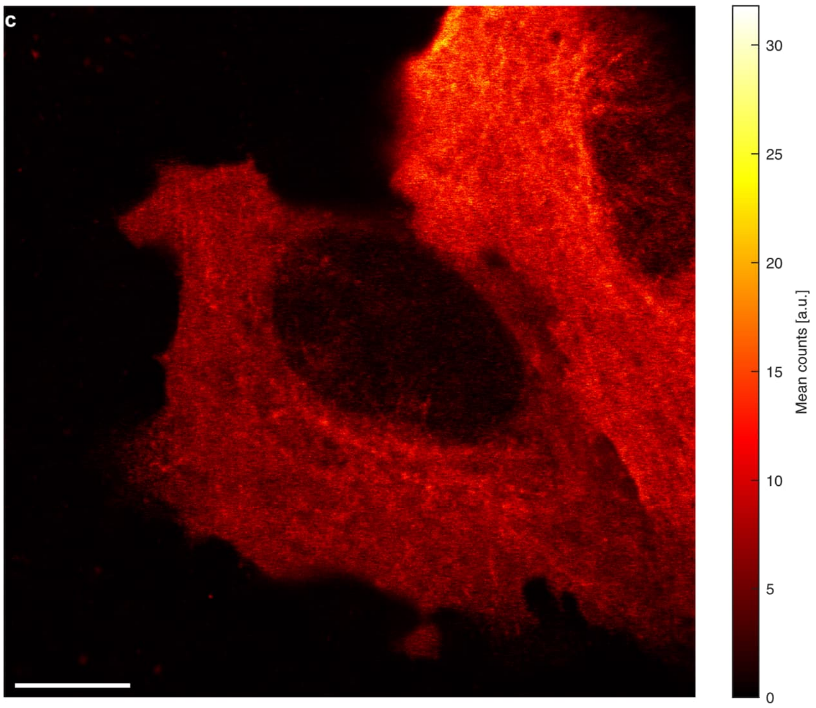


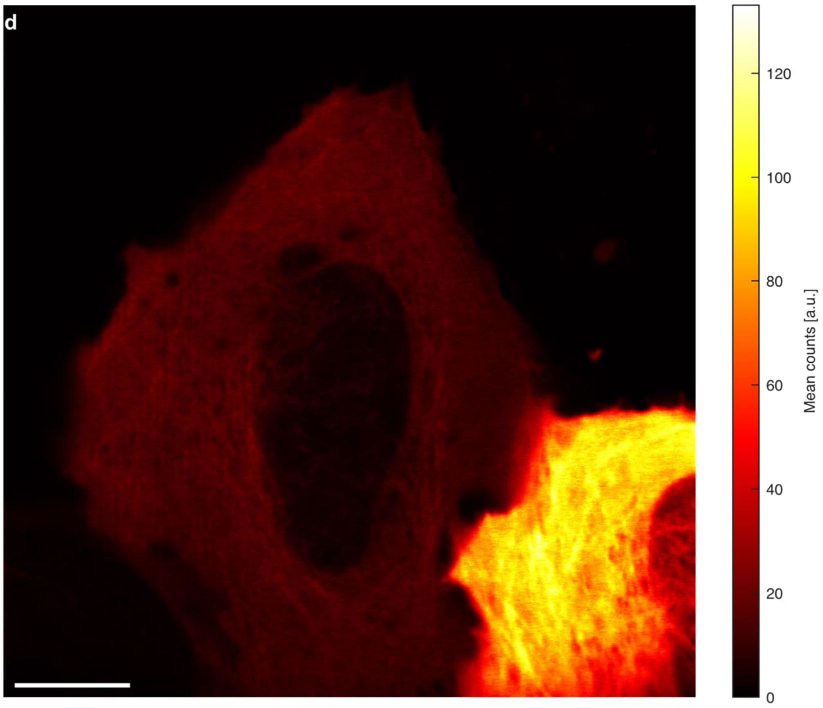


Suppl. Fig. S11 | Confocal overview images of living cells for kinesin tracking. The fluorophore of the dye JF549 was directly bound to the N-terminus of the kinesin. Scalebars 10 µm.

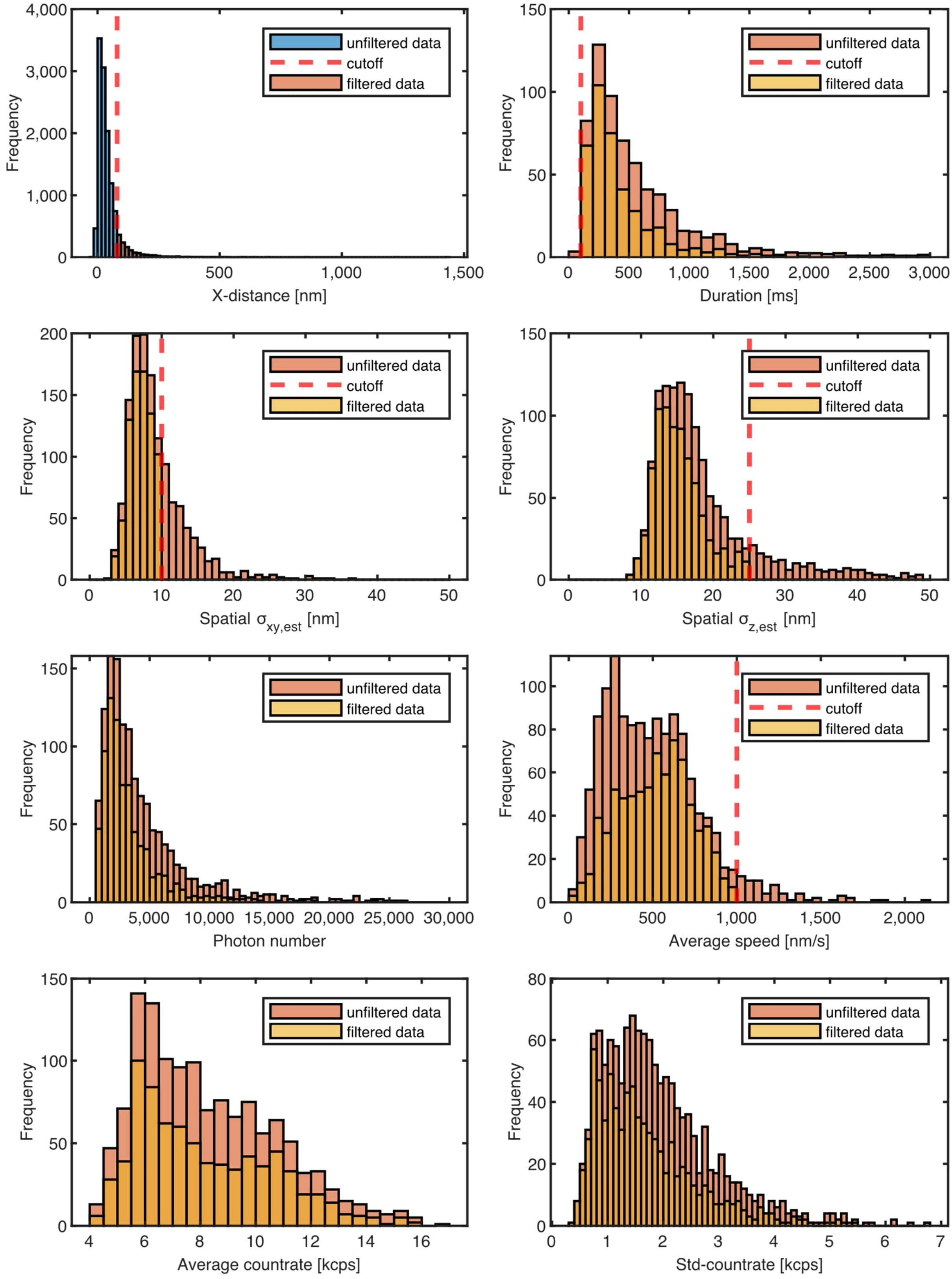


Suppl. Fig. S12 | Characteristic of fluorescence tracks measured by tracking of kinesin in living cells. Histograms of unfiltered and filtered tracks of kinesin tracking in living cells with filter parameters listed in Supplementary Table T2. First, the tracks were filtered by the distance along the principal axis (x-axis) and the retained tracks were filtered by the other filter parameters.

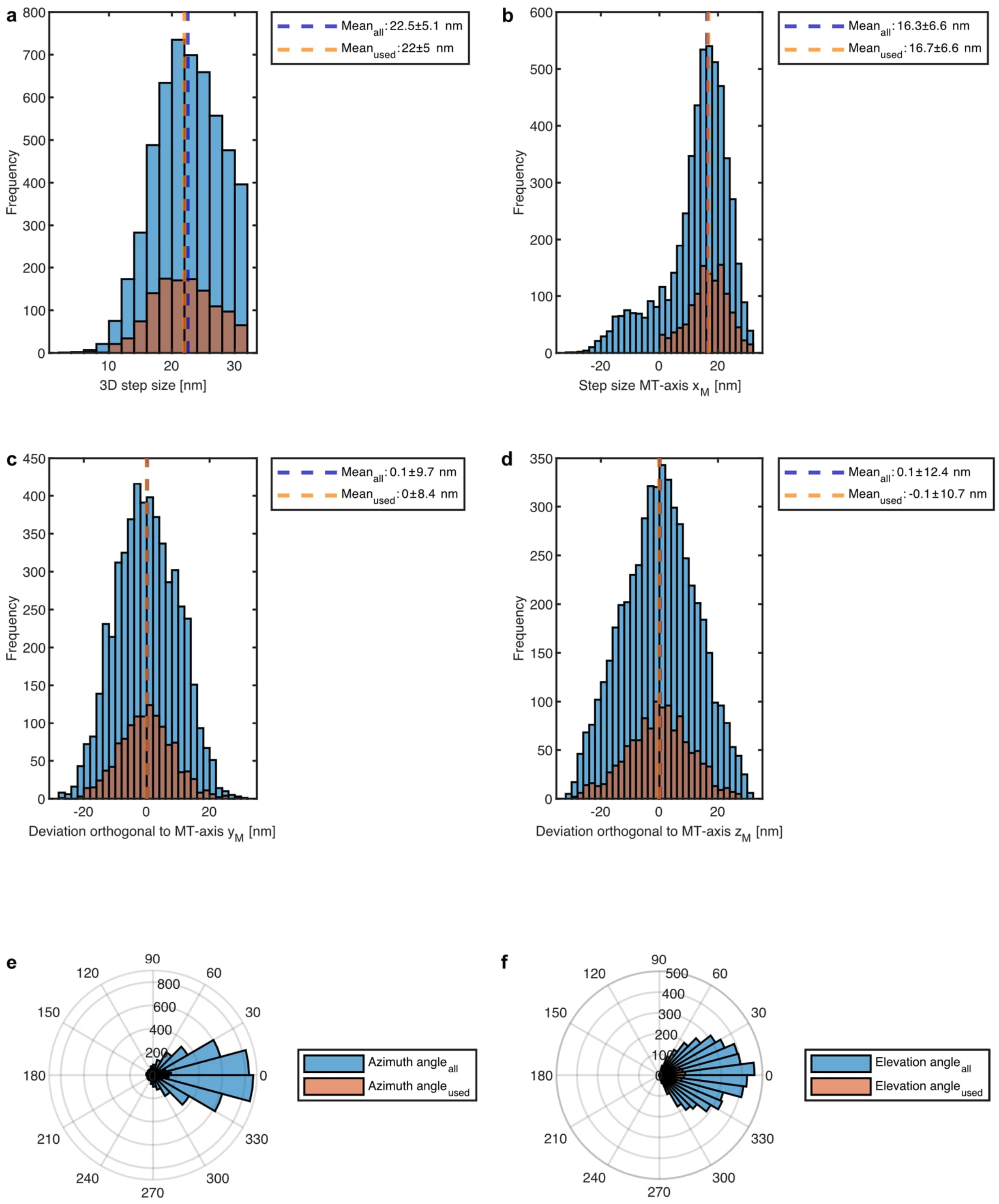


Suppl. Fig. S13 | Characteristics of detected steps in kinesin tracks of living cells. Histograms and mean values of all steps and of retained steps in kinesin tracks established in living cells with filter parameters listed in Supplementary Table T3. (a) Euclidean step size distribution in 3D. (b) Step size along the principal axis and (c,d) the deviation along the orthogonal axis. (e,f) Distribution of azimuth and elevation angle.

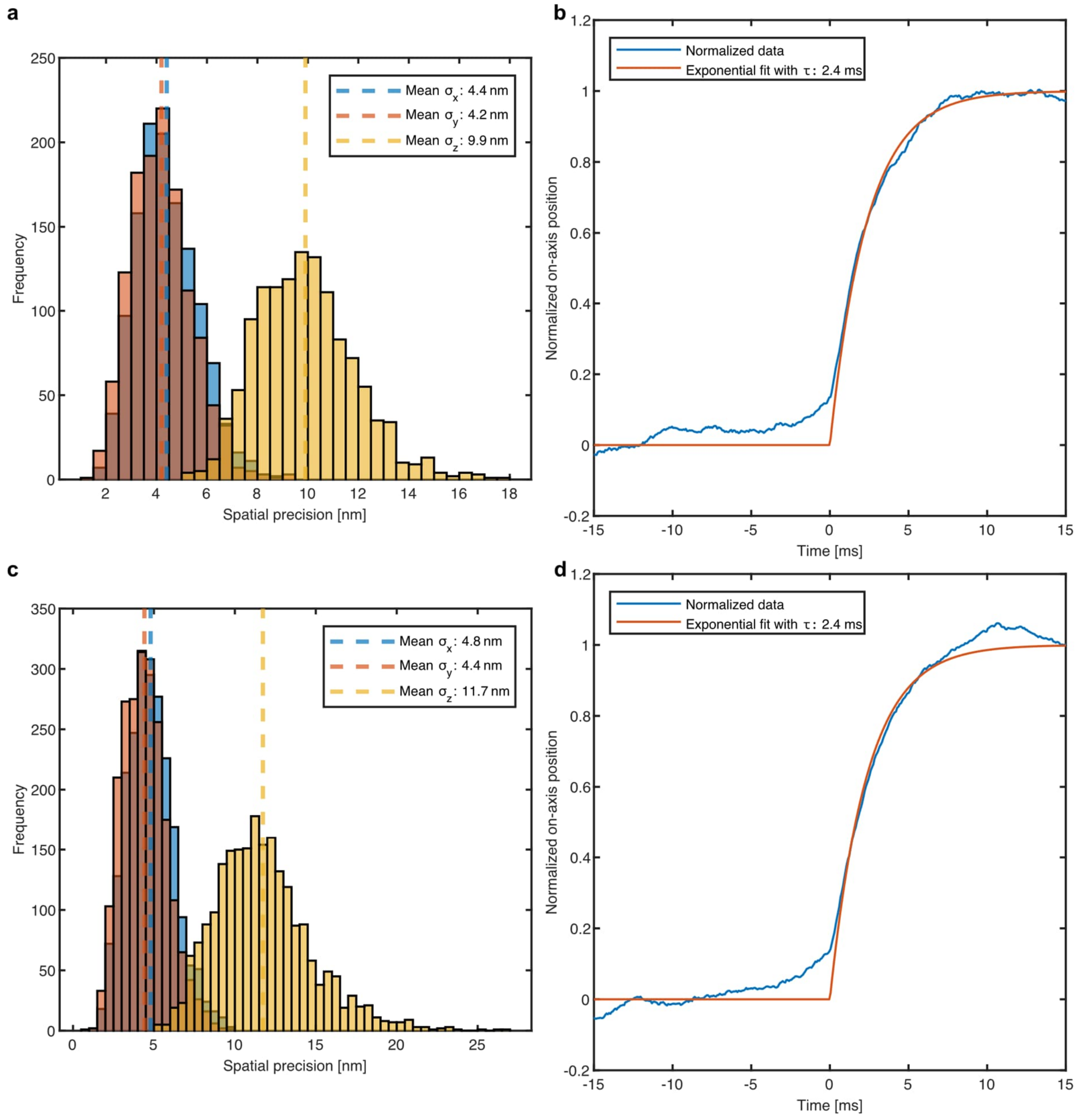


Suppl. Fig. S14 | Spatial and temporal resolution for tracking in fixed and living cells. Spatial resolution histogram and temporal resolution fit of detected center plateaus for tracking in fixed (a,b) and living (c,d) cells.

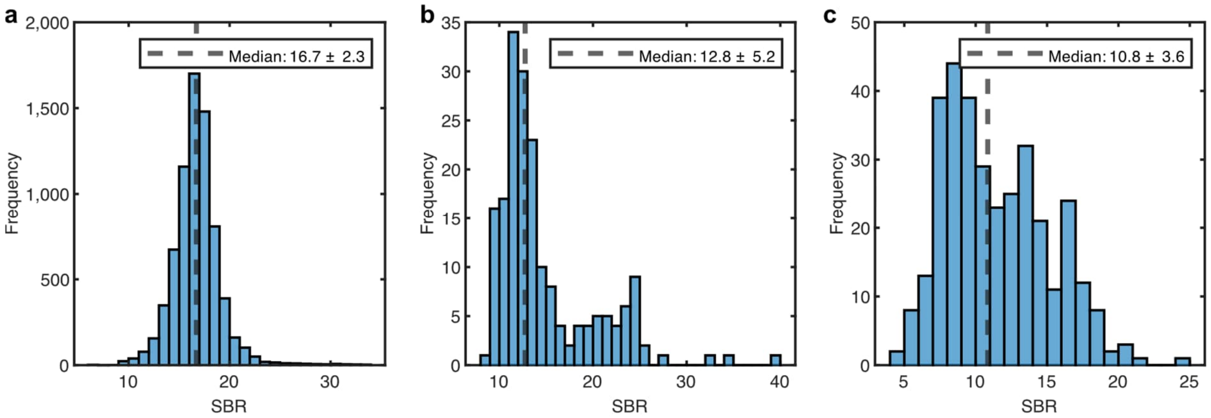


Suppl. Fig. S15 | Signal-to-background ratio histograms for all measured samples. SBR histograms for 3×3 12 nm DNA origami grid (a), tracking in fixed cells (b) and tracking in living cells (c) with their corresponding median and standard deviations.
