## Supplementary figures and images for "3D-MINSTED nanoscopy and protein tracking in densely labelled living cells"

### Supplementary Video 1

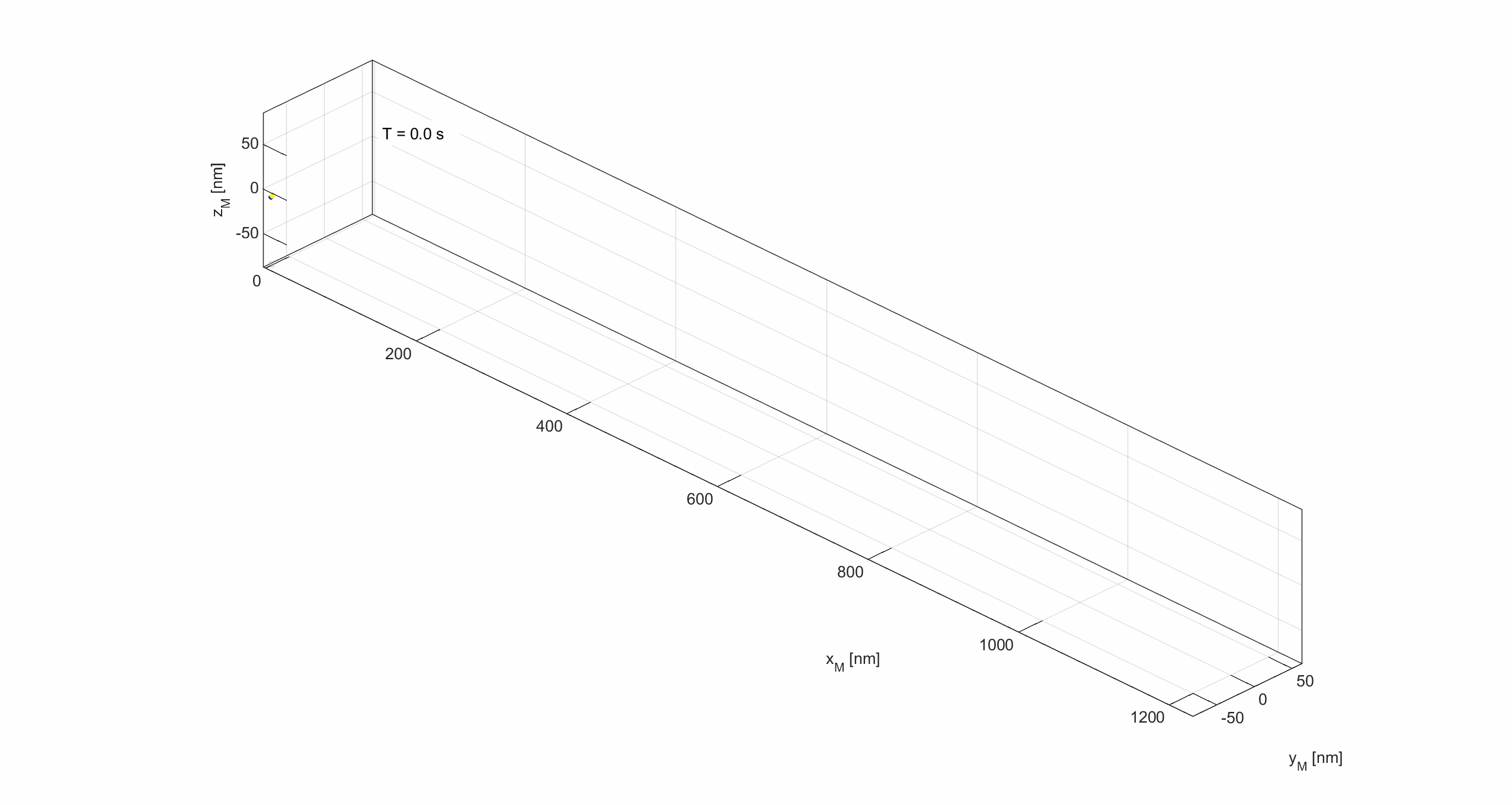
